# Cardiac dysfunction affects eye development and vision by reducing supply of lipids in fish

**DOI:** 10.1101/2021.05.11.443591

**Authors:** Elin Sørhus, Sonnich Meier, Carey E. Donald, Tomasz Furmanek, Rolf B. Edvardsen, Kai K. Lie

## Abstract

Developing organisms are especially vulnerable to environmental stressors. We aimed to understand the underlying mechanisms of phenanthrene (Phe) and crude oil induced eye malformations. We exposed Atlantic haddock (*Melanogrammus aeglefinus*) embryos to a known L-type calcium channel blocker, nicardipine hydrochloride (Nic), and compared to early embryonic crude oil (Oil) and late embryonic Phe toxicity. All treatments lead to severe, eye, jaw and spinal deformities at early larval stages. At 3 days post hatching, larvae from all treatments and corresponding controls were dissected. Eyes, trunk, head and yolk sac were subjected to lipid profiling, and eyes were also subjected to transcriptomic profiling. Changes in lipid profiles and the transcriptome suggested that the dysfunctional and abnormal eyes in our treatments were due to both disruption of signaling pathways and insufficient supply of essential fatty acids and other nutrients form the yolk.

## Introduction

Developing organisms are particularly vulnerable to environmental stressors, such as oil pollution. Early life stages of fish are especially susceptible due to their small size, critical developmental window, limited migration capacities, high lipid content and less developed metabolic pathways. It is well known that crude oil exposure causes severe functional and morphometric symptoms in fish embryo and larval stages^1-3^. In particular, functional and developmental heart abnormalities, jaw and abnormally shaped eyes and/or reduced eye size have been documented^1-7^. Later in life (juveniles and adult stages), these impairments may lead to increased injuries, developmental delays, changes in swimming capacity and mortality^8-10^.

Untangling of the underlying mechanisms for these crude oil induced effects has been a focus of research for decades. The heart has been identified as one of the most impacted organs by crude oil exposure, resulting in both morphological and functional phenotypes^1,11^. Cardiac function can be directly affected by both crude oil and single polycyclic aromatic hydrocarbons (PAHs), such as phenanthrene (Phe). Both crude oil and Phe interfere specifically with the excitation contraction coupling in cardiomyocytes by disrupting rectifier potassium currents and calcium cycling^12-15^. Exposure to Phe increases intracellular calcium in rat and zebrafish embryonic cardiac myoblasts^16^. In addition to its role in the heart, calcium is also a crucial second messenger that is involved in numerous biological functions, like transcription of genes^17^. Accordingly, the calcium regulated genes were induced by crude oil exposure prior to effects on cardiac function, suggesting that crude oil has a direct effect on calcium homeostasis^5^. Although this direct effect on cardiac function is reversible^18^, a transient effect on ion homeostasis may disrupt cardiac development and cause irreversible changes in a developing animal^6^. Similarly, a transient disruption of craniofacial muscular function could also irreversibly affect the outgrowth of jaw structures^19^.

Nicardipine hydrochloride (Nic) is a peripheral vasodilator pharmaceutical drug used to treat hypertension, chronic angina pectoris, and Prinzmetal’s variant angina^20^. Nic inhibits influx of extra cellular calcium across the membrane, inhibits ion-control gating mechanisms and/or interferes with the release of calcium from internal stores like sarcoplasmic reticulum^20^. The inhibitory action for the calcium channel is limited to the voltage-dependent calcium influxes, and does not affect the receptor operated and NaCl-free-induced calcium influxes^21^. Thus, by decreasing intracellular calcium, Nic facilitates the relaxation of the muscle cell^22^.

Exposure to crude oil and oil components have resulted in reduced eye size (anapthalmia) and reduced size with abnormalities (microthalmia) in fish larvae^5,7,23^. Eye development may be affected through several pathways. Vitamin A (retinol) and its biologically active metabolite retinoic acid (RA) plays multiple roles during embryonic eye development^24^. Several environmental contaminants, including crude oil and PAHs, have been shown to disrupt RA signaling either through Cyp induction or other Ahr independent pathways^7,25-27^. Circulatory disorders in early life stages of fish also reduce the eye size^28^, suggesting that supply of essential building blocks and nutrients from the yolk is essential for proper eye development. Sufficient amount of the long-chain fatty acids docosahexanoic acids (DHA, 22:6 (n-3)) have been shown to be critical for the developing eye and good visual performance in early larvae stages of marine fish^29-31^.

In this study we aimed to better understand if the underlying mechanisms of the eye abnormalities found after Phe and crude oil exposure were related to heart dysfunction and reduced circulation of essential lipids. Therefore, we investigated the consequences of a known L-type channel blocker, Nic, and compared to early embryonic crude oil and late embryonic Phe toxicity in Atlantic haddock (*Melanogrammus aeglefinus*). Disruption of developmental signaling pathways in early embryonic stages may cause irreversible impacts on early cardiac development and disrupt circulation at later larvae stages, which we tested with crude oil exposures. Disruption of cardiac function after completion of main organogenesis may also affect circulatory dependent proliferation and formation events^5^, which we tested with Phe exposures. We suspected that circulatory disorders may impact lipid distribution in the organism. Therefore, we collected and dissected yolk sac larvae exposed to crude oil (Oil) and Nic in early embryonic development and larvae exposed to Phe in late embryonic stages. We measured the distribution of lipids in eyes, yolk sac, head and trunk of the yolk sac larvae and mRNA sequenced the eyes of an additional set of larvae from each treatment. The aim was to link abnormal eye development to disrupted signaling pathways and circulation abnormalities.

## Results

Overall, exposure to crude Oil, Phe and Nic resulted in jaw and spinal deformities and smaller and malformed eyes. No ventricular contraction was observed in either Phe or Nic treatment. At 3 dph, changes in fatty acid profile and total cholesterol were found in the eyes of all treatments. Among most affected pathways in the eye transcriptomes were signaling pathways (e.g. *calcium signaling*), muscle function and formation pathways, cholesterol biosynthesis pathways, fatty acid metabolism pathways and phototransduction. For graphical summary of findings, see Supplementary Fig. 1.

### Exposure

Levels of total PAHs in the Oil treatment were approximately 800 µg PAH/L in water, and approximately 1300 µg PAH/g in tissue at exposure stop. Levels of Phe in the water in the Phe treatment were approximately 200 µg Phe/L. Detailed results are in Supplementary Table 1 and Supplementary Data 1 and 2.

### Morphological and functional phenotypes

Body axis deformities at 3 dph were detected in all treatments. Hunched spine was observed in approximately 50-60 % of the Phe treated animals (Fig. 1E), while arched spines and spinal curvatures were seen in all Nic (Fig. 1F) and 53 % of the Oil treated animals (Fig. 1B). Similarly, outgrowth of the jaws was affected in all treatments. While a homogenous pool of animals with similar jaws were found in the Nic treatment (Fig. 1F’), Phe treatment showed animals with jaw deformities similar to Nic animals (Fig. 1E’) and animals with less severe jaw phenotypes (Fig. 1D’). The jaw phenotypes in Oil treatment were severe and more undefinable in many of the animals (Fig. 1B’and 1C’) but did not resemble observations in Phe and Nic treatment.

**Figure 1:**
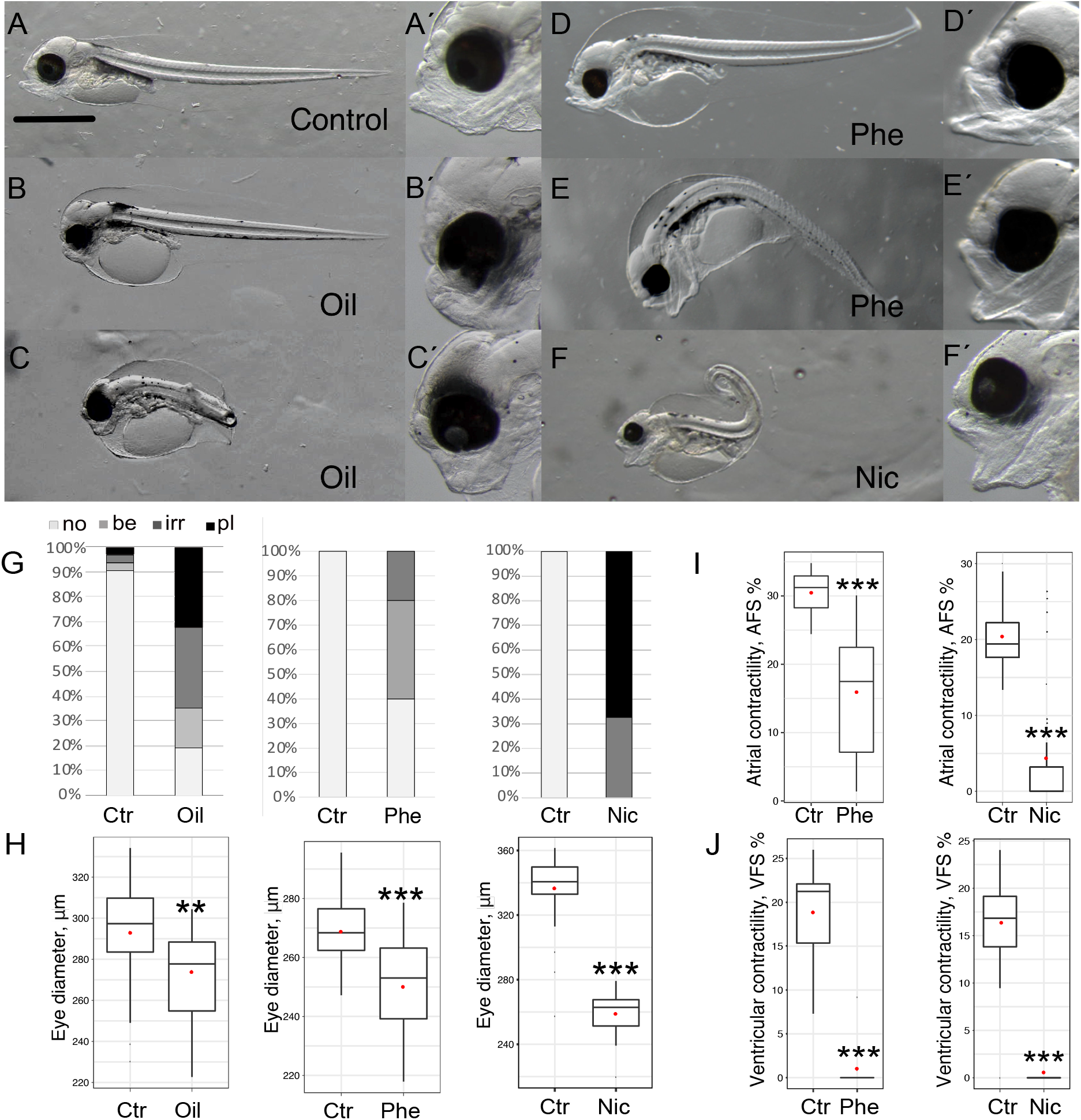
Phenotypes after exposure to crude oil (Oil), phenanthrene (Phe) and nicardipine hydrochloride (Nic). A) Control; B) and C) phenotype after Oil exposure; D) and E) phenotype after Phe exposure; and F) phenotype after Nic exposure. A’) no (no) eye phenotype; D’) bend (be) eye phenotype; E’) irregular (irr) eye phenotype; B’,C’,F’) protruding lens (pl) phenotype. G) Eye phenotype at 3 dph in 30 (Oil and Nic) and 5 (Phe) individuals. H) Eye diameter at 11.5 dpf (Phe) and 3 dph (Nic and Oil) in µm in 30 individuals. I) Atrial and ventricular contractility or atrial/ventricular fractional shortening (AFS/VFS) in % for Phe and Nic exposure (30 individuals). Contractility data for Oil treatment were not available.

The epithelium of the eye appeared to be thinner in all treated animals opposed to the control animals, and often ruptured during dissection. Abnormal eye phenotypes were seen in all treatment. Images of the eye phenotypes normal, bend, irregular and protruding lens are documented in Fig. 1A’-1F’. The most severe phenotype, protruding lens (pl), was only observed in Oil (32%) and Nic (62%) treatments (Fig. 1G). Eye diameter was significantly reduced in all treatments (Fig. 1H), and the reduction was largest in the Nic treatment.

Exposure to Phe acutely affected cardiac contraction and little to no activity was observed during exposure. After 72 hours of exposure (11.5 dpf, Fig. 9) the atrial activity was variable but significantly decreased, while no ventricular contractility was detected (Fig 1I). Similarly on the seventh exposure day at 3 dph, very little cardiac activity and no ventricular contraction was observed. After exposure, the eggs were transferred to clean water. However, Nic irreversibly binds to L-type calcium channel and therefore blocking occurred up to the sampling point at 3 dph. Very little or no cardiac activity was observed after the expected onset of first heartbeat (6.5 dpf) one day after transfer to clean water until 3 dph (Fig. 1I and 1J). Contractility data for the Oil treatment were not available.

### Distribution of lipids in 3 dph larvae

The distribution of lipids and lipid profiles were affected in Oil, Phe and Nic treatments (Table 1, Fig. 2, Supplementary Tables 2-4). Total fatty acids and cholesterol were decreased in the eyes both in total amount pr larvae but also after normalizing to the eye diameter (Supplementary Table 5). In the eyes of all treatments, the FA profile (% of total FAs) showed higher relative amount of saturated- and monounsaturated fatty acids (ΣSFA and ΣMUFA) while polysaturated fatty acids (ΣPUFA) were lower (Figure 2A, B and C). Especially, 16:0, 18:0, 18:1 (n-9) and 22:6 (n-3) fatty acids were highly affected in the eyes (Supplementary Tables 2-4, Fig. 2D, E, F). The relative 22:6 (n-3) amount were highest in the eyes (38-39 % of total FAs in the control fish) compared with the other three compartments (22:6 (n-3)=29-32% (Head), 29-30 % (trunk), 24-29 % (yolk) of total FAs). In the control fish, the ratio between 22:6 (n-3)/20:5 (n-3) was 4.4 in the eyes, while the head, trunk and yolk had much lower ratios, 2.9, 2.7, 2.5, respectively (Supplementary Fig. 2). In the eyes of the treatment groups the ratio between 22:6 (n-3)/20:5 (n-3) was lower compared to control; Oil (3.6), Phe (3.4) and Nic (2.7). In addition, only minor differences were detected between the compartments. In the head, trunk and yolk sac, there were no difference in levels of ΣSFA and ΣMUFA in any of the treatments (Supplementary Tables 2-4). Levels of ΣPUFA and especially 22:6 (n-3) were lower in the heads of the Oil and Nic animals, but not in the Phe animals. Similarly, higher content of 22:6 (n-3) in the yolk was only observed in Oil and Nic treated animals (Fig. 2D, E, F, Supplementary Tables 2-4). The lipid samples clustered according to tissue (eyes, head, trunk and yolk sac), except for Nic and Oil eyes (Supplementary Fig. 3). Detailed fatty acid profiles in eyes, head, trunk and yolk sac for the treatments and respective controls are given in Supplementary Table 2 (Oil), 3 (Phe) and 4 (Nic).

**Table 1.** Amount of fatty acids and cholesterol (µg/larva) in eyes, head, trunk and yolk from control and treatments groups. C, control; Phe, phenanthrene; Nic, Nicardipine. Single measurements were performed on pools of 10 animals (N=1).

| | Total fatty acids ($\mu\text{g/larva}$) | | | | Cholesterol ($\mu\text{g/larva}$) | | | |
| --- | --- | --- | --- | --- | --- | --- | --- | --- |
|  | Eyes | Head | Trunk | Yolk | Eyes | Head | Trunk | Yolk |
| Control | 0.79 | 2.93 | 2.47 | 1.66 | 0.12 | 0.51 | 0.35 | 0.07 |
| Oil | 0.28 | 1.37 | 2.23 | 1.82 | 0.04 | 0.23 | 0.31 | 0.08 |
| Oil/C (%) | 36% | 47% | 91% | 109% | 37% | 45% | 89% | 110% |
| Control | 0.84 | 2.32 | 2.78 | 0.68 | 0.12 | 0.42 | 0.43 | 0.04 |
| Phe | 0.60 | 2.32 | 2.70 | 0.60 | 0.09 | 0.40 | 0.39 | 0.03 |
| Phe/C (%) | 72% | 100% | 97% | 89% | 75% | 94% | 91% | 88% |
| Control | 0.660 | 3.05 | 2.93 | 1.630 | 0.100 | 0.540 | 0.480 | 0.086 |
| Nic | 0.45 | 1.95 | 3.150 | 2.010 | 0.066 | 0.340 | 0.490 | 0.112 |
| Nic/C (%) | 68% | 64% | 108% | 123% | 69% | 63% | 102% | 130% |

**Figure 2.**
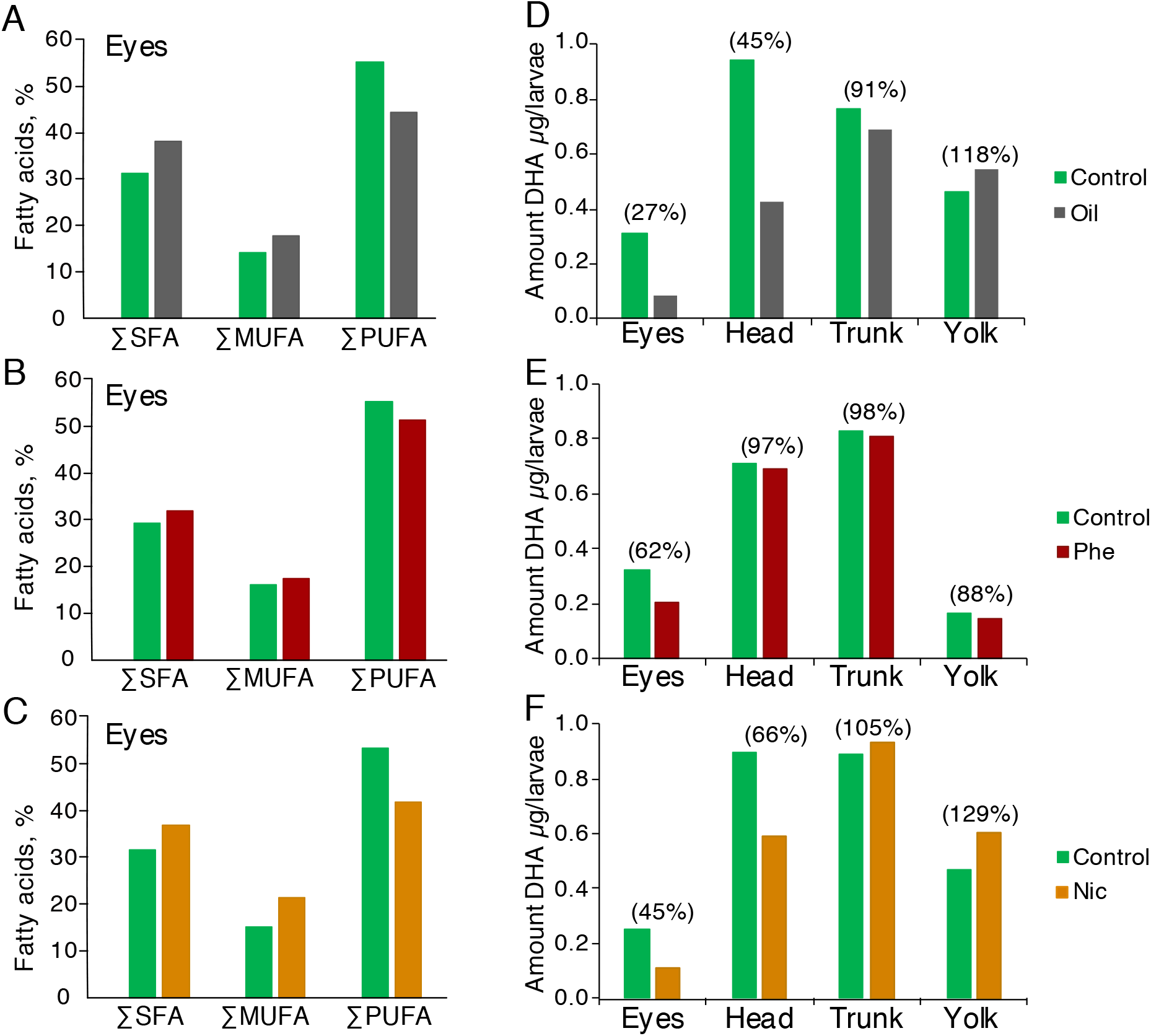
Distribution of fatty acids. Relative content of saturated, mono- and polysaturated fatty acids in eyes of A) crude oil (Oil), B) phenanthrene (Phe) and C) nicardipine hydrochloride (Nic) treatments. Amount of docosahexaenoic acid (DHA) (22:6 (n-3)) in eyes, head, trunk and yolk from control and treatments groups; D) Oil, E) Phe, F) Nic. Single measurements were performed on pools of 10 animals. The number in the bracket represents the amount of DHA in the treatment groups relative to their controls (in %).

### Differentially expressed genes and most enriched pathways in the eyes of 3 dph larvae

Time period and length of exposure and homogeneity in the treatment pools were reflected in number of differentially expressed genes (DEGs) and enriched pathways. Total number of DEGs (p<0.05) with more than 10 transcripts in the Oil, Phe and Nic were 179, 2643 and 11980, respectively. Exposure in the Oil treatment ended 10 days before dissection, and correspondingly, fewer pathways were affected in the Oil eyes. The direction of regulation in the Oil eyes mainly followed Nic eyes which were also exposed during early embryogenesis (Fig. 9). IPA Canonical Pathways and Diseases and Biofunctions with activation z-score threshold of 3 or more are presented in Fig. 3A and B, while Fig. 3C displays common pathways among top 40 KEGG pathways in Phe and Nic eyes. For top 40 relative KEGG pathways for all treatments, see Supplementary Fig. 4. In general, signaling pathways were activated in the Phe eyes, and inhibited in the Nic eyes (Fig. 3A, Supplementary Fig. 5). Contrarily, IPA *Cholesterol biosynthesis* (Fig. 3A), IPA *Fatty acid metabolism* (Fig. 3B), *Synthesis of lipid* (Fig. 3B), *Metabolism of membrane lipid derivative* (Fig. 3B) and KEGG *Steroid biosynthesis* (Fig. 3C) pathways were activated in both Phe and Nic eyes. Oil eyes also showed an up-regulation of genes in KEGG *Fatty acid biosynthesis* pathway (Supplementary Fig. 4A) in addition to *Transport of lipid* (Fig 3B). Phototransduction pathway were one of the few other KEGG pathways common for all treatments (Supplementary Fig. 4). Down-regulation was found in Oil and Nic eyes, while mainly an up-regulation was observed in the Phe eyes. Likewise, *Retinal degeneration* (Fig. 3B) was activated in Nic while inhibited in Phe eyes.

**Figure 3:**
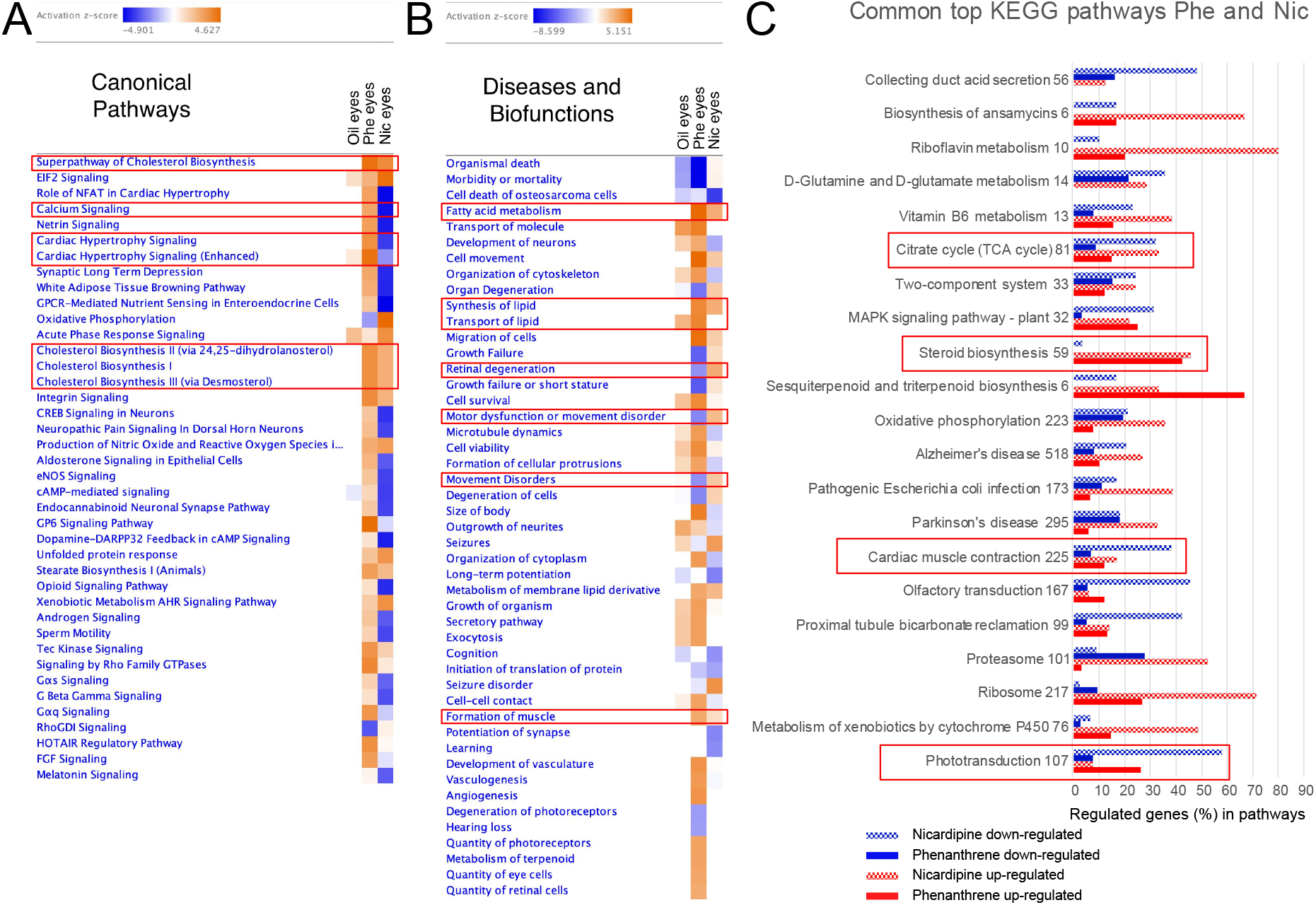
Top affected IPA and KEGG pathways in eye tissues. Overview of most enriched IPA Canonical pathways (A) and IPA Disease and Biofunctions (B), z-score threshold was set to 3 and p-value threshold was set to 0.05. The figure is displayed with z-score. Orange: pathway activated. Blue: pathway inhibited. Darker colors indicate stronger activation/inhibiton. C) Common relative top KEGG pathways for Phe and Nic eyes (Oil eyes were not included due to low number of DEGs). Fold change threshold was set to 1.5 and p-value to <0.05. Number of genes regulated in pathways was divided by number of total annotated genes pathway and are presented as percentage on x-axis. Number behind the Pathway name represents total number of annotated genes in respective pathway. Red and blue bars represent up-regulation and down-regulation, respectively. Outlined pathways: pathways related to fatty acid and cholesterol metabolism, muscle function and formation, calcium signaling and eye development and function. IPA; Ingenuity Pathway Analysis . KEGG; Kyoto Encyclopedia of Genes and Genomes. Oil; crude oil treatment, Phe; phenanthrene treatment, Nic; nicardipine hydrochloride treatment.

### Fatty acid and cholesterol homeostasis

*Fatty acid metabolism* was among top IPA Diseases and Biofunctions and was activated in both Phe and Nic eyes (Fig. 3B). In the same way, the KEGG pathways *Fatty acid biosynthesis, degradation, elongation, fat digestion and absorption* and *PPAR signaling* were highly regulated, and mainly up-regulated (Supplementary Dataset 3). Several DEGs were shared between *PPAR signaling* and *Fatty acid biosynthesis* and *degradation*. All or most DEGs were exclusive for *Fatty acid elongation* and *Fat digestion and absorption* (Supplementary Dataset 3.). Several DEGs involved in the above-mentioned pathways were common for Phe and Nic eyes (Supplementary Dataset 3). Among these DEGs were Fatty acid desaturase 2 (*fads2)* and Fatty acid elongase 6 (*elovl6)* which were up-regulated 2.9 and 3.8 (*fads2)* and 3.2 and 2.9 fold (*evovl6*) in Phe and Nic eyes, respectively (Supplementary Dataset 3). Another *elovl, elovl4*, were highly expressed in the eyes regardless of treatment (e.g. *elovl4* (IMR10009452) with approximately 7000 normalized transcripts, Supplementary Dataset 3). However, expression levels were reduced in the Nic treatment (FC = -2.5, 3000 normalized mean transcripts, Supplementary Dataset 3). Upstream regulator indicated reduced levels of high-density lipoproteins (HDL), cholesterol, lysophosphatidylcholine and DHA in Nic and Phe exposed eyes, while PPARGC1b and PPARD were increased in the same samples (Supplementary Fig. 6).

Cholesterol pathways were also activated in both Phe and Nic eyes (Fig. 3). A condensed pathway combining *Mevalonate pathway* and *Superpathway of cholesterol* show that almost all genes involved in the pathways were up-regulated (Fig. 4). Interestingly, 3-hydroxy-3-methylglutaryl-CoA reductase (*hmgcr*) and 3-hydroxy-3-methylglutaryl-CoA synthase 1 (*hmgcs1*) were only up-regulated in the Phe eyes. Another interesting gene up-regulated was the TLC domain containing protein 1 (*tlcd1*) which was up-regulated in both Phe and Nic treatment.

**Figure 4:**
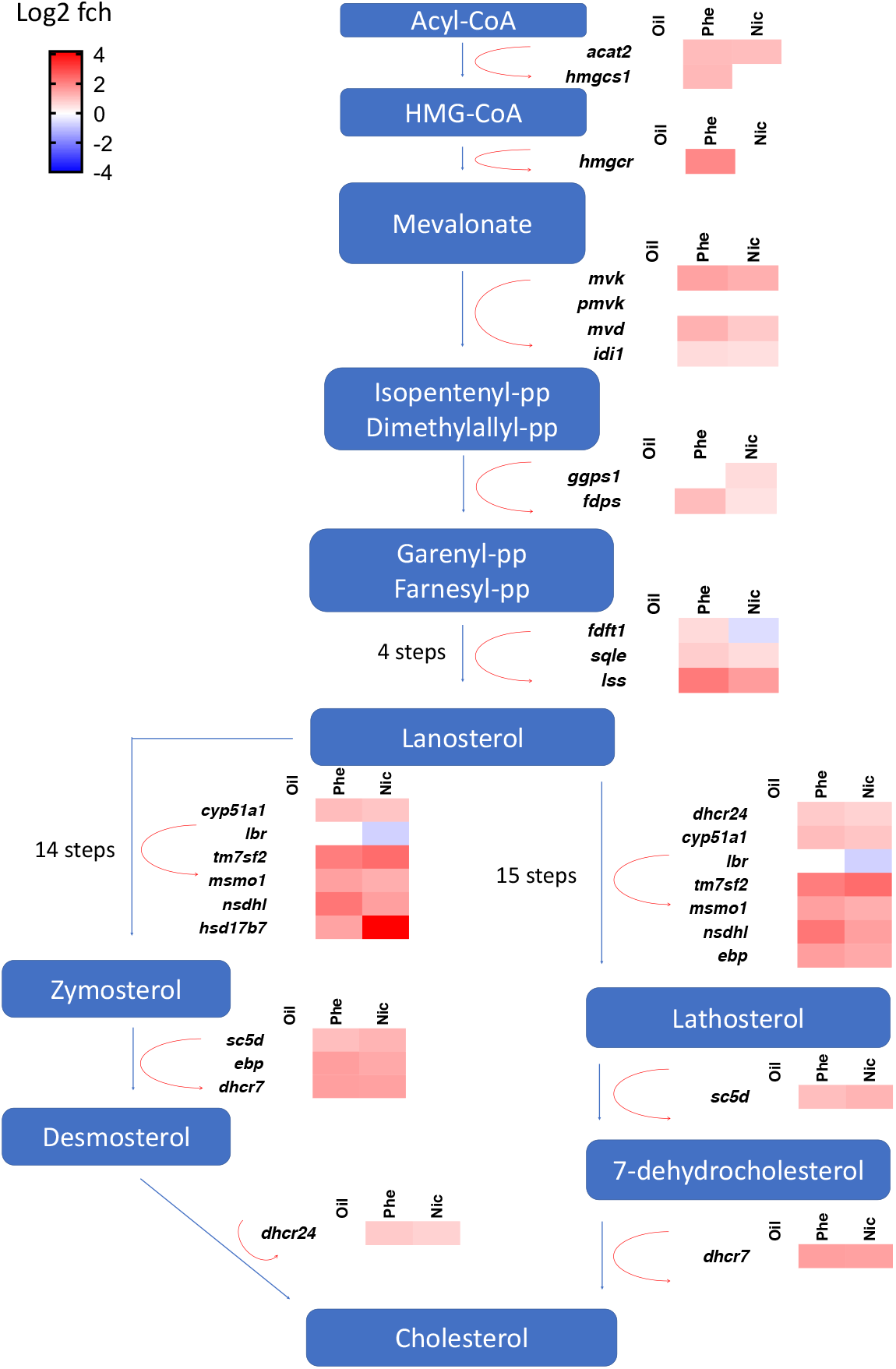
Superpathway of cholesterol and Mevalonate pathway. Mevalonate pathway and Superpathway of cholesterol combined. Differentially expressed genes indicated as heatmaps next to the respective step. Oil; crude oil treatment, Phe; phenanthrene treatment, Nic; nicardipine hydrochloride treatment, Log2FCH; base 2 logarithm of the fold change.

### Calcium signaling and muscular function

Calcium signaling pathway was among IPA Canonical pathways with highest activation score (Fig. 3A). Only one DEG (*grin1*) was common for all treatments, while we found 28 common DEGs for Phe and Nic eyes (Fig. 5A), but mainly with opposite expression (Fig. 5B). For example, the genes encoding L-type channels (*cacna1f*), skeletal muscle Ryanodine receptor 1 (*ryr1*) and inositol 1,4,5-trisphosphate receptor type 1 (*itpr1*) were up-regulated in Phe eyes and down-regulated in Nic (Fig. 5 and Supplementary Data 4). Oil eyes only had 6 DEGs in *Calcium signaling* pathway, all common to and in the same direction as Nic eyes (Fig. 5C).

Similar to *Calcium signaling* pathway, the Disease and Biofunctions *Motor dysfunction or movement* and *Movement disorders* were oppositely activated in Phe vs Nic. Here, Nic was activated and Phe was inhibited (Fig. 6). A visual web of the DEGs (262) in *Motor dysfunction and movement disorder* in Phe eyes overlapping with the Canonical pathways (CP), *Calcium signaling* (21 genes) and *Cardiac hypertrophy signaling (Enhanced)* (19 genes) are shown in Fig. 6A. A complementary web of the same genes for Nic eyes (Supplementary Fig. 7B) mainly shows the opposite expression. Similarly, KEGG *Cardiac contraction* pathway, which in this context reflects effect on muscular function in the eyes and not cardiac contraction, was also oppositely regulated in the two treatments (Supplementary Fig. 8, Supplementary Data 5). *Formation of muscle* on the contrary, was activated in both treatments (Fig. 6A). To visualize common DEGs in between some of these pathways, Venn diagrams including IPAs *Calcium signaling, Cardiac hypertrophy signaling, Motor dysfunction and movement disorders* and *Formation of muscle* were made and are displayed in Fig. 6B. Especially *Calcium signaling* pathway had several DEGs in common with the other pathways. For example, in the Nic eyes, only 19 of 108 DEGs were exclusive for *Calcium signaling* pathway.

**Figure 5:**
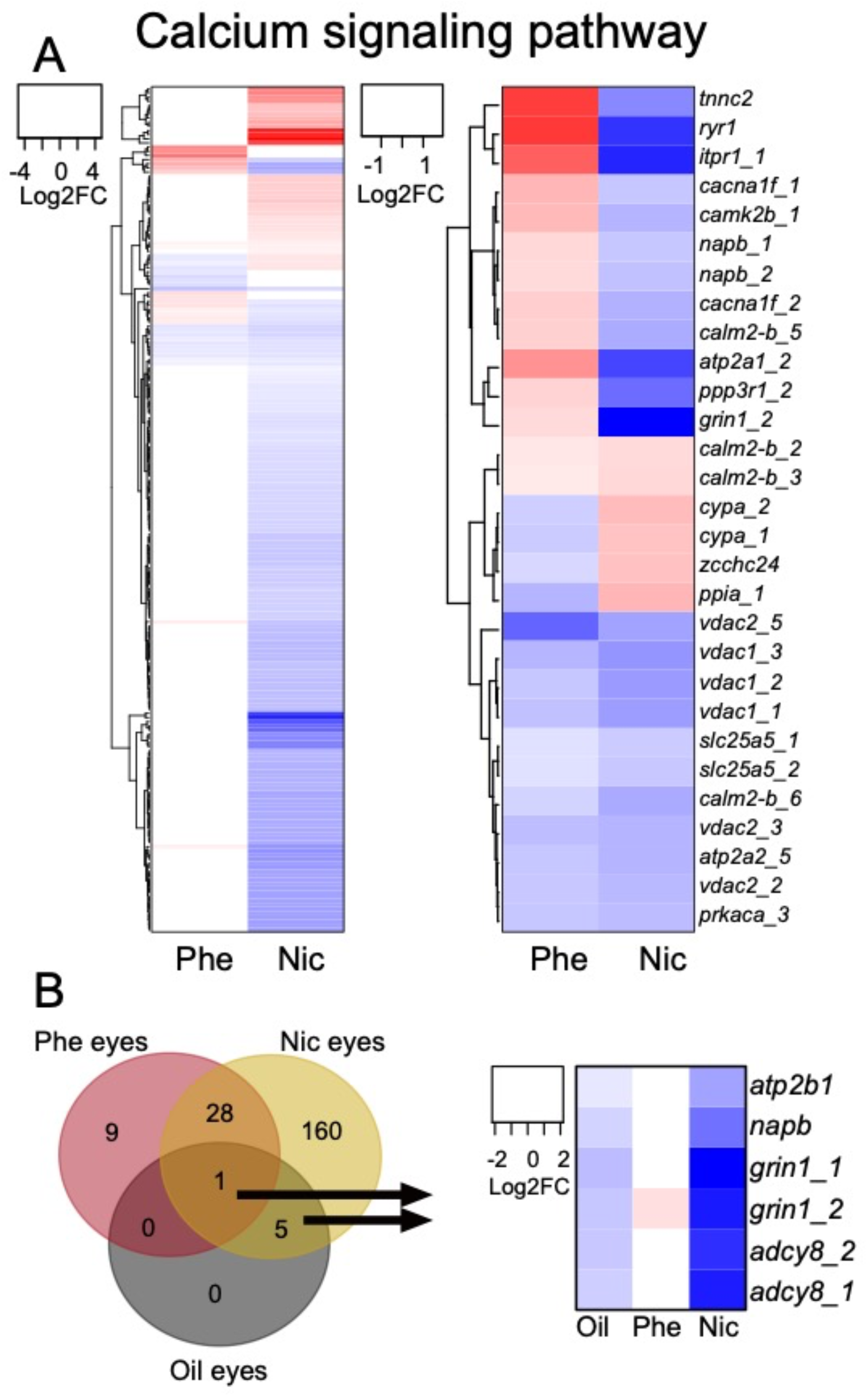
Calcium signaling pathway. A) KEGG Calcium signaling pathway. Left: All differentially expressed genes (DEGs) in genes in Oil, Phe and/or Nic eyes. Right: DEGs common for Phe and Nic eyes. B) Left: Venn diagram for DEGs in KEGG calcium pathway for all treatments. Right: Heatmap of all DEGs in Oil eyes within KEGG Calcium signaling pathway. Oil; crude oil treatment, Phe; phenanthrene treatment, Nic; nicardipine hydrochloride treatment.

**Figure 6:**
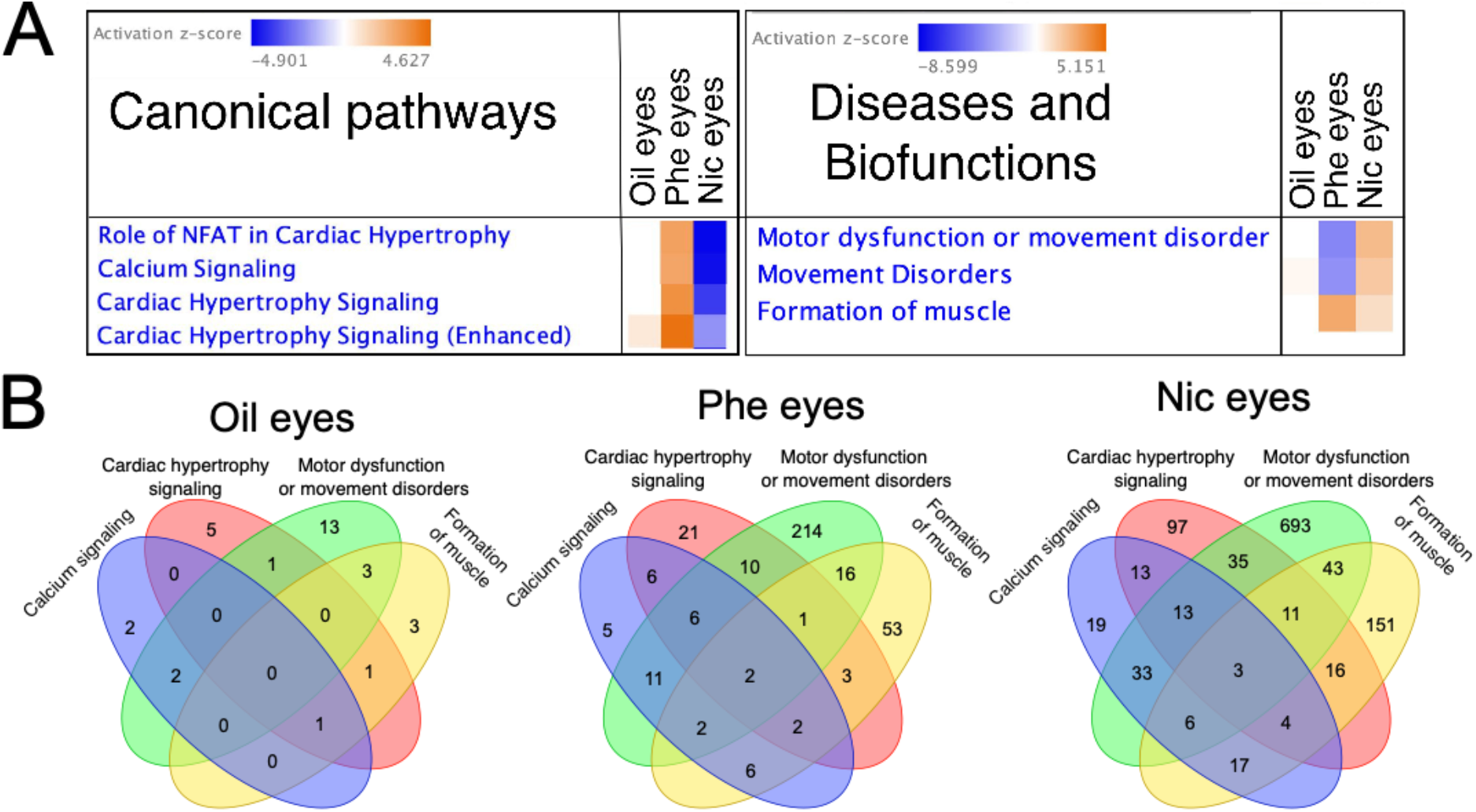
Calcium signaling pathway, motor dysfunctions and muscle formation. A) IPA z-score tables for selected Canonical pathways and Diseases and Biofunctions. B) Venn diagram showing common differentially expressed genes in IPA Calcium signaling pathway, Cardiac hypertrophy signaling (Enhanced) pathway, and Motor dysfunction or movement disorders and Formation of muscle. Threshold p-value for DEGs >0.05. Oil; crude oil treatment, Phe; phenanthrene treatment, Nic; nicardipine hydrochloride treatment.

### Retinol metabolism and Phototransduction

Retinol metabolism was affected in Phe and Nic eyes but not in Oil eyes. Phototransduction pathway, on the other hand, was affected in eyes for all treatments (Fig. 7).

In *Retinol metabolism*, ten of the DEGs were common for the Phe and Nic treatments (Fig. 7A). Genes were mainly up-regulated in Phe eyes, while more variable in the Nic eyes (Fig. 7B). In *Phototransduction* pathway most genes were down-regulated in Nic eyes, while more variable in Phe eyes. Oil eyes had only 12 regulated genes which all were common with findings in Nic eyes (Fig. 7A) and also showed the same direction as Nic eyes (Fig. 7B). Phe and Nic eyes had 30 genes in common (Fig. 7A), and only 8 were regulated in the same direction (Fig. 7B).

**Figure 7:**
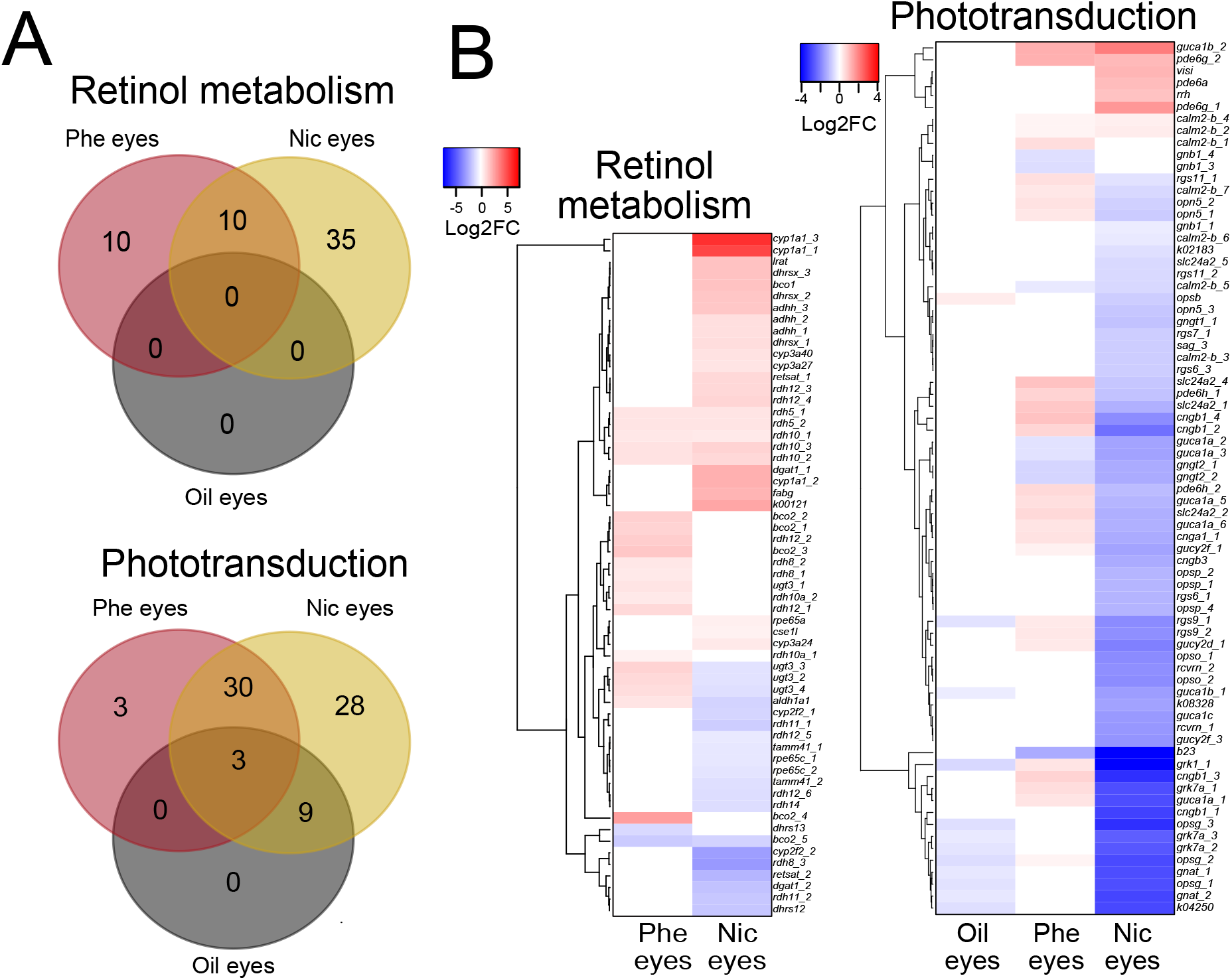
Retinol metabolism and phototransduction. A) Upper: Venn diagram of differentially expressed genes (DEGs) in Retinol metabolism pathway in Phenanthrene eyes (Phe eyes), Nicardipine eyes (Nic eyes) and crude oil exposed whole embryos at 6 days post fertilization (dpf) (Oil whole 6 dpf). Lower: Venn diagram of DEGs in Phototransduction pathway in Phe eyes, Nic eyes and Oil eyes at 3 dph. B) Heatmap of DEGs for the various treatments in Retinol metabolism pathway and Phototransduction. Gene with ID K00121 was not annotated to genebank but corresponds to ADHH, Alcohol dehydrogenase class 3 chain H in KEGG. Phe; phenanthrene treatment, Nic; Nicardipine hydrochloride treatment, Log2FC; base 2 logarithm of the fold change.

## Discussion

The present study was designed to identify underlying mechanisms causing the eye malformations following crude oil exposure previously observed^2,5,7^. Exposure of embryos/larvae to Oil, Phe and Nic caused severe abnormalities visible at 3 dph. Lipid distribution was also disrupted, which was linked to transcriptomic changes in pathways related to cholesterol and lipid homeostasis. Signaling pathways, especially calcium signaling, were affected in Phe and Nic treated animals, however the activation was opposite, i.e. suggesting an opposite effect on calcium homeostasis in the two treatments. In comparison, few genes were regulated at 3 dph in the Oil treatment, which was 10 days after end of exposure. Both inappropriate signaling and deficit of essential nutrients may lead to the observed eye malformations observed in all our treatments.

The initial intention for the Nic treatment was to expose in the same developmental period as the Oil exposure (i.e 2.5 -5-5 dpf). However, Nic continues to block cardiac function after transfer to clean water, and we suspected that the exposure was irreversible. Nic is readily metabolized in the liver with a half-life of 8.5 hours in humans^20^. But there is no knowledge on the metabolization rate in the early life stages of cold-water fish species. The haddock embryos do not have fully developed liver until open mouth stage (1 dph)^32^, with the rudimental liver cells visible at 7 dpf^33^ and liver bud appearing approximately at hatch^2^. Thus, an incomplete metabolism of the drug is likely, resulting in a partly irreversible exposure in the developmental period examined. To emulate the effect that Nic was having at later larval stages, we performed the Phe exposure.

The consequences of an exposure are dependent on the exposure length and the developmental period of exposure^2,6^. In present study, we found that the Nic treatment gave the most homologous severely malformed population. The morphological phenotypes in the Phe treatment were more variable. However, all animals showed the same functional phenotype, a non-contracting or silent ventricle that demonstrated the acute effect Phe has on cardiac function^13,18^. In the Oil treatment, morphological phenotypes were also variable but more severe, reflecting various impacts from multiple toxicants in the complex oil mixture during an essential period of development (2.5 dpf-5.5 dpf). Also, the Oil treatment would be expected to suffer from prolonged exposure due to oil droplet fouling on the eggshell^34,35^, but the exposure definitely ended at hatching^2,6^.

Spinal and jaw deformities could be secondary to cardiac dysfunction^28^, a result of signal disturbance (mainly in early embryonic development)^36,37^, or a direct effect on muscular function^19,38^. After 72 hours of Phe exposure, no ventricular contraction was observed, suggesting a specific impact on cardiac ventricular function. Accordingly, tricyclic PAHs such as Phe are known to impact cardiac function directly^28^ by disturbing potassium and calcium currents essential for excitation contraction coupling (ECC) in the cardiomyocytes^13-15^. Nic blocks the L-type calcium channel and inhibits influx of calcium across the membrane of both myocardial and smooth cells^22^ and skeletal muscle cells^39^, although the predominant effect is on arterial smooth muscle cells in humans^22^. We expected an impact on all muscle fibers in the Nic treatment. However, it is not known whether Phe’s impact on ECC is solely reserved for cardiac tissues or would have the same effect in all excitable tissues like with the Nic treatment. Appropriate outgrowth of jaws is dependent on craniofacial muscle function^19^, and therefore paralyzing craniofacial muscles would result in similar abnormalities to what we observe in our treatments. In addition, Nic and Oil animals were exposed during organogenesis, potentially affecting development of craniofacial elements^40^. Interestingly, despite that Phe was exposed in late organogenesis, the Nic craniofacial phenotypes resemble the Phe phenotypes more than Oil phenotypes. This suggests that opposed to Oil treatment, Nic exposure do not seem to affect patterning of jaw but only affect outgrowth of the jaws.

There is a close interdependent development of the spinal cord and muscle fibers along the cord^41^. A direct effect on skeletal muscle fibers could potentially result in the spinal abnormalities seen in our treatment. The difference in phenotype, i.e. arching spine in Oil and Nic and hunched spine in Phe treatment, could either be a product of timing of exposure or slight differences in mechanistic effect on muscular function. Similar to Nic treatment, silent heart morpholino zebrafish also possess severe jaw and spinal disorders^28^. This supports that jaw and spinal deformities are secondary to circulation, since the gene knocked out in silent heart morpholino zebrafish is the cardiac specific troponin 2 (*tnnt2*), necessary for cardiac contraction. In comparison, Nic blocks L-type channel in cardiac, smooth and skeletal muscle cells^22^. Thus, Nic induced disruption of skeletal muscle function is likely to cause spinal and craniofacial abnormalities. In summary, the observed jaw and spinal deformities found in our treatment are most likely due to multiple impacts including direct effect on skeletal muscle, and not only secondary to circulation disorders.

The fatty acid and cholesterol analysis suggested a disrupted transport of lipids from the yolk to the eye with possible implications on development. The altered lipid distribution was also reflected in the transcriptomic data, indicating activation of *Fatty acid metabolism* and *Cholesterol biosynthesis* pathways in the exposed eyes. Lipids are essential for eye development and growth^29,30^. In line with other studies, n-3 PUFA, DHA (22:6 (n-3)), and the saturated fatty acids, stearic acid (18:0) and palmitic acid (16:0) accounted for the largest proportion of fatty acids^31^. PUFAs contribute to the physical chemical properties of the cell membrane by influencing neurogenesis, neuroplasiticy, neurite growth, synaptogenesis and membrane permeability and fluidity^42^.

Long chain (LC)-PUFAs cannot be synthesized *de novo* in vertebrates and must be consumed or obtained from yolk sac either intact or from a select group of precursors such as α -linolenic acid (18:3n-3) and linoleic acid (18:2n-6)^43^. We observed reduction of PUFAs and especially DHA in the eyes of all treatments, a finding that was supported by the predicted decrease of the up-stream regulators DHA and HDL following IPA analysis. We also observed increased levels of SFAs and MUFAs, mainly palmitic acid, stearic acid and oleic acid. The changes in fatty acid profile and genes related to lipid metabolism suggests that the eyes are insufficiently trying to compensate for the deprivation of essential fatty acids by *de novo* synthesis of SFA and MUFAs and synthesis of LC-PUFAs by increasing desaturation and elongation of precursors. (Fig. 8).

**Figure 8.**
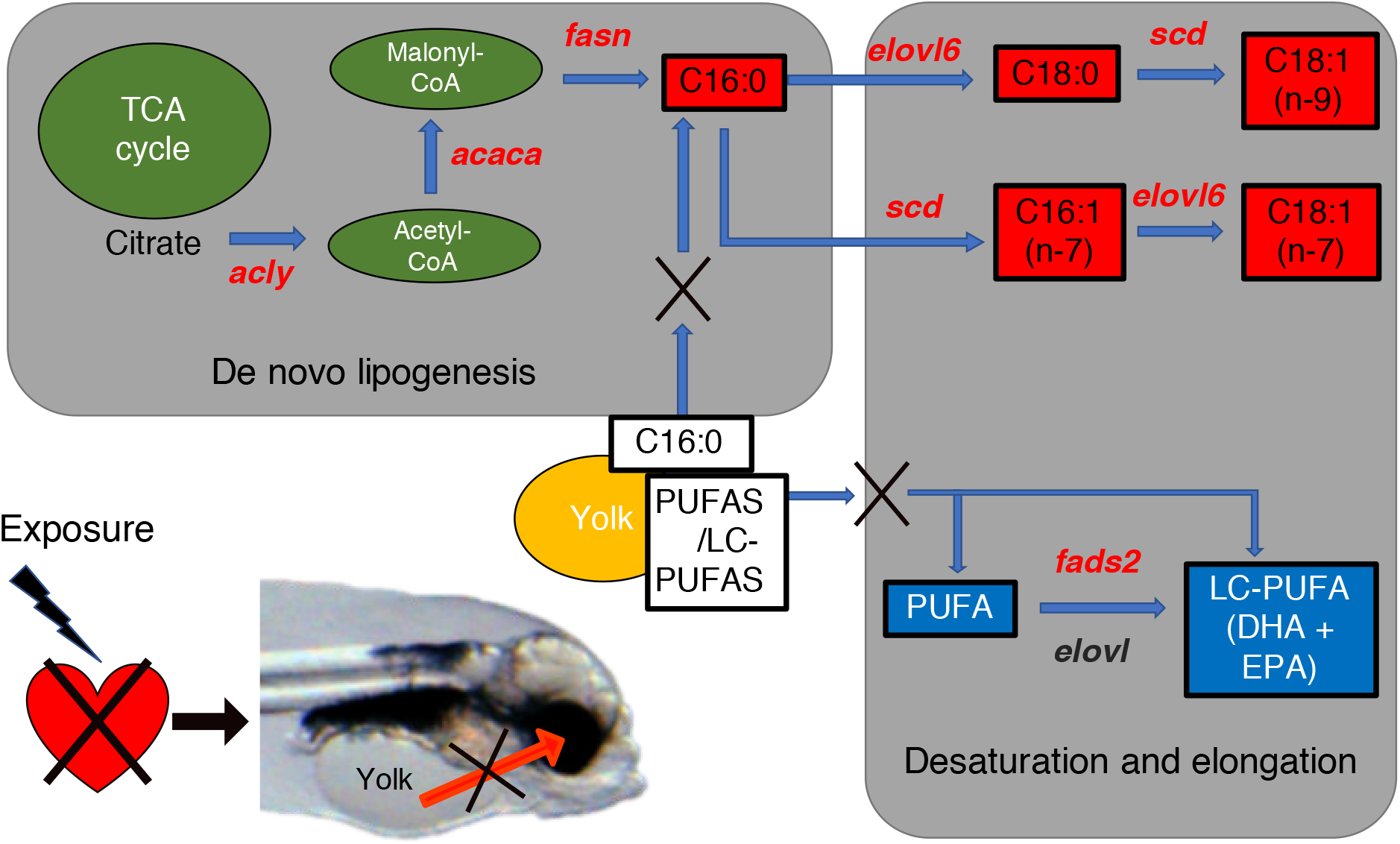
Schematic overview of circulatory related effect on lipid metabolism. The data suggest that cardiac dysfunction following exposure disrupts transport of lipids from the yolk to the eye during embryonic development. As a result, increased expression of lipid metabolizing genes (red letters) in the eye facilitates de novo synthesis of saturated fatty acids (red boxes). In addition, decrease in allocation of LC-PUFAS from the yolk induces the main desaturase gene (fads2) in an effort to compensate the lack of DHA in the eye. Red and blue boxes indicate observed increase and decrease in specific fatty acids following chemical exposure (Phe and Nic).

Very long chain fatty acids (VLC-PUFA C_24_-C_38_) play important roles in eye development of fish larvae^44^. The levels of these VLC-PUFAs in the organism are low, and therefore demand sensitive detection methods^45^. In present study we used conventional gas chromatography with flame ionization detector (GC-FID) and were only able to detect up to C_24_ PUFAs. But the elongase of VLC-PUFAs;C_24_, Elovl4, which are known to be highly expressed in the eyes and brain^44,46^, were also abundant in our eye-transcriptomes. However, *elovl4* were not up-regulated in our chemically treated groups, on the contrary two *elovl4* paralogs were down-regulated in the Nic treatment. These findings suggests that the larvae were not able to increase the synthesis of VLC-PUFAs to compensate for the reduced amount of PUFA (especially DHA) coming from the yolk.

PUFAs and cholesterol are essential for maintaining the integrity and fluidity in cell membranes^42,47^. A dynamic membrane composition is necessary for optimal function in varying conditions. One of the major mechanisms of maintaining membrane fluidity under changing conditions is through regulating the fatty acid groups in the membrane and cholesterol content. For example, in colder environments, enrichment of unsaturated fatty acids, such as DHA, was observed^48^. Incorporation of lipophilic oil components into the membrane are proposed to impact the fluidity of the membrane and possibly alter membrane protein functions^49^. A recent study showed that pretreatment with cholesterol partially mitigated the effect of Phe on heart rate^50^. Although most genes in the cholesterol biosynthesis pathway were upregulated by both Nic and Phe, the rate limiting enzymes, 3-hydroxy-3-methylglutaryl-CoA reductase (*hmgcr*) and 3-hydroxy-3-methylglutaryl-CoA Synthase 1 (*hmgcs1*) were only up-regulated in Phe eyes. How Phe disrupt/block these channels is still unknown. In vitro studies show that Phe has the ability to incorporate into the cell. Likewise, alkyl phenols and estrogen altered the fatty acid profile to contain more SFA and less n-3 PUFA in cod (*Gadus morhua*), the changes in lipid profile in our treatments could also indicate a compensatory response to altered membrane fluidity. Accordingly, we observed up-regulation of a protein that regulates fluidity of the plasma membrane (*tlcd1*). A plausible mode of action for disruption of potassium and calcium channels could be alteration of the channel by disturbing the fluidity around the channel.

To summarize the effects regarding lipids in the eyes, we observed a decrease of total fatty acids, cholesterol, and PUFAs. With this, we observed increased activation of fatty acid metabolism and cholesterol biosynthesis. All these effects could be due to a) a circulation-dependent deprivation of lipids and cholesterol, and b) compensatory response to increased cell membrane fluidity.

Enriched pathways in 3 dph eyes of Phe and Nic treatment included signaling pathways and muscular function pathways, confirming the consequences of an impact on essential ion currents in muscle cells of the eyes. However, opposite z-activation score in the two treatments suggests an opposite cellular effect. Nic reduces cytoplasmic calcium by reducing influx of extracellular calcium, but also inhibits ion control gating mechanisms and possibly interfere with release of calcium from sarcoplasmic reticulum^20,22^. An opposite z-activation score was observed in the Phe treatment, suggesting that Phe increases intracellular calcium. Accordingly, increased intracellular calcium was also observed in rat embryonic cardiac myoblasts (H9C2) exposed to Phe^16^. The regulation of the gene *itpr1* is indicative of the opposite effect on calcium homeostasis in Nic and Phe treatment. Itpr1 controls the calcium induced calcium release (CICR) of calcium from the endoplasmic reticulum (ER) to the cytoplasm^51^. Increase of calcium in the cytoplasm induces release of more calcium to the cytoplasm from internal storages such as ER. Accordingly, we observed up-regulation of *itpr1* in Phe eyes and down-regulation in Nic.

Several of the genes involved in *Calcium signaling* pathway were also represented in *Motor dysfunction or movement disorders* and *Formation of muscle* in the various treatments suggesting detection of down-stream effects on muscular function and development in the treatments. Particularly in the Oil treatment, we mainly expected to detect gene expression linked to the down-stream effects of an oil exposure during early embryonic development. For example, only two genes were exclusive for *Calcium signaling*, while 5, 13 and 3 were exclusive for *Cardiac hypertrophy signaling, Motor dysfunction or movement disorders* and *Formation of muscle*, respectively. Overall, we suggest that an impact on calcium homeostasis induced muscular dysfunction and abnormal eye muscle development in the treatments. Regardless of the direction of impact on signaling pathways in Phe and Nic eyes, the consequences for muscle development were similar.

Our data suggest that the animals treated with Oil or Nic during early development resulted in blindness or reduced vision. In contrast, treatment with Phe in late embryonic development resulted in a more compensatory response. In Phe eyes, eye development and function were likely affected by deprivation of essential nutrients and disruption of retinol metabolism. The retina of the eye is highly enriched with LC-PUFAs^52^, and are essential for eye development. PUFAs inhibit the expression of fatty acid desaturase 2 (*fads2*) and elongation of long-chain fatty acids family member 6 (*elovl6*)^53,54^. An up-regulation of *fads2* and *elovl6* indicate reduced amount of PUFAs in Phe and Nic eyes. Accordingly, less n-3 PUFAs (DHA) were observed in the eyes of all treatments compared to control. Due to their critical function, PUFAs in the yolk are reserved for development and not consumed for energy^55^. However, with circulatory defects in Phe and Nic eyes, the essential PUFAs do not reach their destination and therefore results in impaired eye development.

Retinol, often referred to as vitamin A, is a precursor for a range of biological active metabolites including vision, limb patterning, growth and normal cell homeostasis^56^. The active metabolite retinoic acid is a known teratogen and disruption of retinoid signaling, has been shown to cause developmental abnormalities^57,58^. Increased retinoic acid levels and disrupted retinoid signaling has been linked to eye deformities in crude oil exposed animals^7^. Retinol metabolism in the eyes was affected by both Nic and Phe exposure, no DEGs involved in Retinol metabolism were found in oil eyes 10 days after end of exposure. Transcriptomic changes were evaluated during exposure in a previous study, and oil induced expression of *rdh8* was seen at 6 dpf^7^. Up-regulation of *rdh8* was also observed in the Phe eyes, suggesting acute disturbance of retinol metabolism in the Phe treatment. Abnormal Cytochrome P450 (Cyp)-balance can also disrupt retinoic acid signaling pathway through disturbance of Cyp26, and consequently lead to abnormal eye development^7,59^. Cyp1a is believed to be one of the most important factors disrupting retinoid signaling following toxicant exposure^26,60^, and is the main gene induced in crude oil exposed animals^5^. Another Cyp1 paralog, Cyp1b, is also involved in eye development (regulating ocular fissure closure) through both retinoic acid-dependent and -independent pathways^25,61^ and were thought to partly induce the stage dependent eye abnormalities found in Sørhus et al. 2021^5^. Cyp1b induction could also explain the increased levels of retinoic acid observed in conjunction with eye deformities in our previous study^7^. Phe is known to be a poor inducer of Cyp1a^62^, and Phe eyes did not show any effect on any *cyp*s except *cyp51*. However, Nic eyes showed induced expression of both *cyp1a* and *cyp1b* which therefore could contribute to the observed abnormal eye development in this treatment.

In the phototransduction pathway several functional units found in photoreceptor cells (e.g. *opsg, opsb*)^63^ or genes encoding proteins involved in signal transduction (e.g. *gnat* and *rgs9*)^64^ were down-regulated in Oil and Nic eyes. Phe eyes on the other hand, showed an up-regulation of several genes involved in *Phototransduction*. In haddock embryos, *Synaptic vesicle cycle* were among most regulated pathways between 6 and 8 dpf, while *Phototransduction* was among most regulated pathways between 10 dpf and hatching^65^. We therefore suggest that exposure during early embryonic development (Oil and Nic) affects eye development and results in reduced vision or blindness later in life, while a late embryonic exposure mainly impact phototransduction and therefore results in a compensatory response like the one observed in Phe eyes. In conclusion, we suggest a double impact on eye development and phototransduction in our treatments. Inappropriate cardiac function could lead to inadequate supply of essential fatty acids necessary for proper eye development and function and affect transport of other nutrients, such as retinoids. Additionally, disturbance of signaling in pathways such as *Retinol metabolism* may lead to abnormal eye development.

## Conclusion

Treatment with Oil, Phe and the L-type channel blocker Nic lead to severe eye, jaw and spinal deformities. In addition, cardiac malfunctioning was observed in Phe and Nic treated animals. Nic blocks L-type channel in all muscle tissues. Therefore, the spinal and jaw abnormalities in Nic treated animals were most likely due to disrupted muscular function and not only lack of circulation. The Nic phenotypes resembled phenotypes found in Oil and Phe treatment, thus, we suggest that a direct effect on skeletal muscle function also could be the underlying mechanism of the jaw and spinal abnormalities found in these larvae.

The lipid profile was altered in all three treatments. All treatments showed a decrease of total fatty acids, cholesterol and PUFAs and an increase of SFA and MUFAs in the eyes. Accordingly, the treatments showed high impact on cholesterol biosynthesis and fatty acid metabolism. This observation could reflect both disrupted transport of nutrients from the yolk to the eyes due to dysfunctional circulation and compensatory responses to changes in membrane fluidity. Further, the eye transcriptome of the Phe and Nic suggested an opposite effect on calcium homoeostasis in these treatments. Phe treatment likely resulted in an increase of intracellular calcium, while Nic treatment resulted in a decrease. This in turn led to opposite impact on muscle function, but down-stream effect on muscular development was the same. Our data suggest that dysfunctional and abnormal eyes in our treatments were due to both disruption of signaling and insufficient supply of essential fatty acids and other nutrients form the yolk.

## Acknowledgments

We would like to acknowledge Stig Ove Utskot for breeding and management of the fish, Therese Aase and Arve Fossen for carrying out the lipid analysis and Holly Shiels for discussions of the idea and interpretation of some of the data. This work was financed by the Research Council of Norway (EGGTOX: Unraveling the mechanistic effects of crude oil toxicity during early life stages of cold-water marine teleosts (Project # 267820, www.forskningsradet.no) and the Institute of Marine Research, Norway. The funders had no role in study design, data collection and analysis, decision to publish, or preparation of the manuscript.

## Ethics declarations

### Competing interests

The authors declare no competing interests.

### Author contribution

ES: Conceptualization, Methodology, Investigation, Formal analysis, Data curation, Writing – original draft. CED: Formal analysis, Data curation, Review of paper. SM: Conceptualization, Methodology, Formal analysis, Data curation, Review of paper. TF: Methodology, Resources, Data curation, Review of paper. RBE: Methodology, Resources, Data curation, Review of paper. KKL: Investigation, Formal analysis, Data curation, Writing – original draft.

## Methods

### Passive dosing preparation

A method for silicone passive dosing of Phe (Sigma-Aldrich) was adapted from Vergauwen et al. 2015^66^ and Sørensen et al. 2019^67^. Cords of silicone were precleaned by soaking in methanol overnight followed by two washes of ultrapure water and storage in ultrapure water until use. Approximately 1 g of Phe was added to 30-50 mL of methanol (until saturation) to make the loading solution. Silicone cords were added to the methanol loading solutions and allowed to infuse for two days. After loading, the silicone strips were wiped clean of any solid residues, and methanol was removed by three >2-hour washes in ultrapure water. Three days prior to the exposure, the loaded and rinsed silicone strips were placed in the bottom of a 100 mL glass beaker filled with 60 mL autoclaved seawater.

### Animal collection, management and exposure set up

Fertilized eggs were collected from brood stocks of Atlantic haddock kept at the Austevoll Research Station of the Institute of Marine Research, Norway. Eggs were kept in incubators at 8 ± 2 °C before transfer to exposure tanks, 100 mL beakers or 12-well plates. Experiment overview for all treatments is given in Fig. 9.

**Figure 9:**
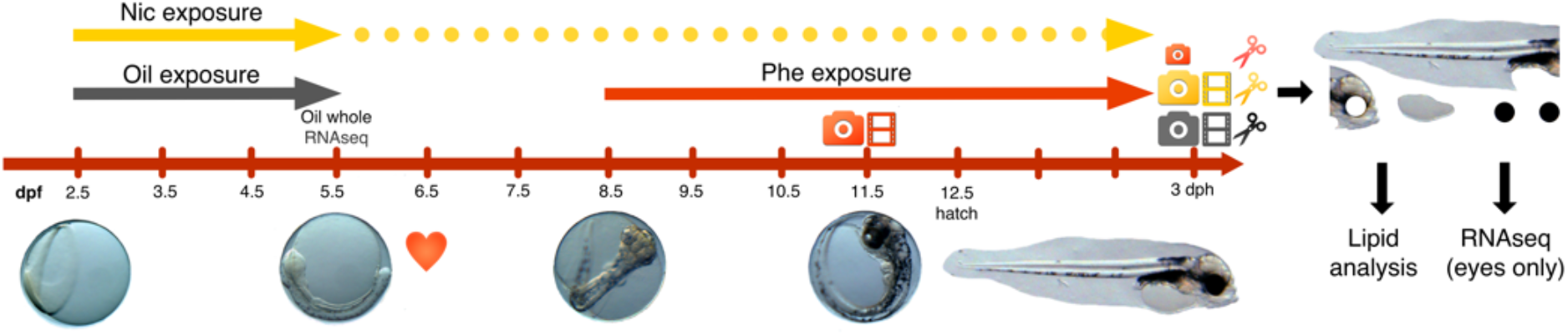
Experimental overview. Exposure period for the crude oil (100 µg oil/L)(Oil) and Nicardipine hydrochloride (Nic) (10 µM) treatment lasted for 72 hours from 2.5 dpf to 5.5 dpf. After exposure the eggs were transferred to clean water, however, Nic binds to a large degree irreversible to L-type calcium channel and therefore the blocking occurs until the sampling point at 3 dph. Phenanthrene (Phe) exposure started at 8.5 dpf to the sampling point at 3 dph. Images and videos were taken at 11.5 dpf for Phe treatment and 3 dph for Oil and Nic. All treatments and associated controls were dissected at 3 dph. Eyes, yolk sac, trunk and head for 10 animals per treatment were subjected for lipid analysis, while eyes from an additional set of animals were subjected for mRNA sequencing. At 5.5 dpf whole embryos in the Oil treatment were sampled and subjected for mRNA sequencing. Heart symbol: Time of first heartbeat.

#### Oil treatment

Approximately 5000 embryos at 2.5 days post fertilization (dpf) were transferred to two 50 L green plastic experimental tanks with flow of 32 L/hr, water temperature of 8.0°C and 12/12 light/dark regime. The crude oil was a weathered blend from the Troll oil field, and the exposure regime was performed as described in Sørhus et al., 2016^2^. The exposure lasted for 72 hours, from 10% epiboly to 10-20 somite stage (2.5-5.5 dpf) (Fig. 9). Experimental setup consisted of one control tank (0 µg oil/L) and one oil tank (100 µg oil/L).

#### Phe treatment

Approximately 100 living eggs were transferred to baked glass exposure jars at 8.5 dpf. One jar contained Phe loaded silicone cord and the control contained cord loaded in clean methanol. Exposure lasted until 3 days post hatching (dph), i.e from two days after first heartbeat to yolk sac larva stage.

#### Nic treatment

Approximately twenty-five embryos at 2.5 dpf were transferred to 6 replicate wells in a 12 well plate containing either 10 mL clean autoclaved saltwater or autoclaved seawater spiked with 10 µM Nicardipine hydrochloride (Cayman Chemical, USA) (n = 6, approx. total number of embryos = 300). The exposure lasted from 2.5 dpf to 5.5 dpf, and the exposure solution was exchanged every day during the exposure. At 5.5 dpf, eggs from all replicates were transferred to a common glass beaker with clean sea water (one beaker for Nic treated animals and one for control). Water was exchanged every second day until end of experiment (3 dph). The Nic experiment was repeated twice to verify consistency, and the replicate experiments resulted in identical morphological and functional phenotypes.

The exposure scenario for Phe with approximately 200 µg Phe/L (1.12 µM) is not environmentally relevant but chosen to induce an effect. Similarly, the relatively high dose of Nic (10 µM) was given to assure appropriate blocking in the treatment group. The Oil treatment on the contrary, was given an environmentally relevant concentration (100 µg oil/L), reflecting a low to medium exposure in an acute oil spill (Sørhus et al., 2015).

### Analytical chemistry

#### Oil exposure

Water samples (1 L) were taken from the exposure tanks at the beginning and the end of the experiment (0 h and 72 h), preserved by acidification (HCl, pH < 2) and stored at 4 °C in the dark until further handling. Details are given in Sørensen et al. 2017^34^. Samples for PAH body burden were collected daily during exposure (24 h, 48 h, 72 h) and one day after exposure (96 h). Pooled samples (50-100 animals) were preserved by flash-freezing in liquid nitrogen and stored at −80 °C. Prior to analysis, tissue samples were extracted by solid-liquid extraction and cleaned-up by solid phase extraction (SPE).

#### Phe exposure

Water samples (1.0 mL) were taken for a replicate study performed in the same period and under exact same conditions (Donald et. al *in prep*). These samples were therefore expected to be representative for the Phe exposure in the current study. Samples were collected from three replicate jars immediately before beginning exposures and each day of a three-day exposure period. The water samples were extracted after addition of internal standard using liquid-liquid extraction with two 1 mL volumes of dichloromethane. Water was removed from the extracts with 0.2-0.4 g Na_2_SO_4_, and extracts were solvent exchanged into isooctane and reduced to 200 µL before analysis on GC-MS/MS.

Agilent 7890 gas chromatograph coupled to an Agilent 7010c triple quadrupole mass detector (GC-MS/MS) was used to perform quantitative analysis of the water samples and body burden. Methods were based on the methods described in Sørensen et al. 2016^68^.

### Morphological and functional measurements

Digital images of un-anaesthetized larvae were immobilized in a glass petri dish containing 3% methylcellulose/97% seawater and positioned on a thermally regulated microscope stage (Brook Industries, Lake Villa, IL) at 8°C. Pictures of 30 individuals were recorded using Moticam 1080 camera (Motic®, Richmond, BC, Canada) mounted on Olympus SZX10 stereomicroscope at 11.5 dpf (Phe) and 3 dph (Nic and Oil treatment). To assess cardiac morphology and function, 20-second videos were captured from 11.5 dpf embryos (Phe) and 3 dph larvae (Nic). Videos were also captured for the Oil treatment, but contractility data from the video analysis of the Oil treatment were rejected because of indications of imaging/sampling related induced stress. Due to low larvae number and no video equipment available at the 3 dph sampling for the Phe experiment, only five representative Phe treated larvae and respective control were photographed at 3 dph using an Olympus SZX10 stereomicroscope equipped with a 5 Mp resolution Infinity camera (Infinity 2–5c, from Lumenera).

Diameter of eye (µm) and ventricular and atrial systolic and diastolic diameters (µm) were processed using imageJ software (Schneider et al., 2012). Atrial and ventricular fractional shortening (FS, %) were then calculated as ((diastole-systole)/diastole)*100. Eye shape phenotypes were described according to the categories: normal (no), bend shape (one indent) (be) irregular shape (two or more indents) (irr) and protruding lens (pl) (see description of eye phenotypes in Sørhus et al. 2021^5^).

### Dissection of 3 dph larvae

Eyes, yolk sac, trunk and head of 3 dph larvae were separated using fine Dumont#55 Forceps (F.S.T. Fine Science Tools) in an Olympus SZX10 stereomicroscope (see Fig. 9). The larvae were anesthetized by adding 500 mg/L MS-222 (Tricaine methanesulfonate, TS 222, Sigma-Aldrich) to a final concentration of 25 mg/L prior to dissection. For RNA-extraction: Three replicates of 10 (Oil and Nic) or 5 (Phe) pair of eyes were either added to RNAlater (Invitrogen) (Oil) or directly to TRIzol reagent (Thermo Fisher Scientific) (Phe and Nic) and kept at -20°C until total RNA extraction. For lipid analysis: 10 pair of eyes, 8-10 yolk sacs, 10 trunks and 10 heads (with eyes excluded) from each treatment and respective controls were pooled and collected in separate 2 ml glass vials (one tube per tissue type per treatment). The samples were snap frozen in liquid Nitrogen and kept at -80°C until analysis. The eye diameter used in Table 1 was measured in another representative set of animals at 3 dph (Oil and Nic) and 11 dpf (Phe).

### Lipid analysis

The different tissue samples (eyes, yolk sac, trunk and heads) were extracted with 3×1 mL with chloroform:methanol (2:1 v/v) and the extract were transferred to 16 ml glass tubes and evaporated to dryness by nitrogen gas, before methylation of all fatty acids to their respective fatty acid methyl esters (FAME). Cholesterol and FAME were analysed on a HP-7890A gas chromatograph (Agilent, USA) with a flame ionization detector (GC-FID) according to a method described in Meier et al. 2016^69^.

Fatty acids (as 51 FAMEs) were identified by comparing retention times with a FAME standard (GLC-463 from Nu-Chek Prep. Elysian, MN, USA). Retention index maps and mass spectral libraries (GC-MS; http://www.chrombox.org/data) were performed under the same chromatographic conditions as the GC-FID^70^.

### Extraction of RNA, sequencing and bioinformatics

Extraction of total RNA from the three eye-pools of 10 pair of 3 dph eyes from Oil, Phe and Nic treatments including respective controls was performed using a combination of Trizol/chloroform extraction and Qiagen’s RNeasy micro column-based RNA extraction. The material was homogenized in 500 µl TRIzol reagent. Thereafter, 100 µl chloroform (Sigma-Aldrich) was added to each tube, and the tubes were shaken vigorously for 2 minutes. The samples were incubated at room temperature for 5 minutes before centrifugation at 10,000xg for 18 minutes at 4°C. After centrifugation the aqueous phase were collected and added to 300 µl 100% ethanol. The samples were transferred to a Qiagen RNeasy Micro column, and the rest of the procedure followed the Qiagen’s instructions which also included a DNase I step. The samples were eluted in 15 µl RNase free water.

cDNA library preparation and sequencing were performed by the Norwegian Sequencing Centre (NSC, Oslo, Norway) using the Illumina strand-specific TruSeq RNA-seq library Preparation Kit. A total of 18 samples were sent to the sequencing facility and 16 were subjected for analysis (two samples were lost during cDNA library preparation): Three (Oil/control, Phe/control) and two (Nic/control) biological replicates. Paired-end libraries were sequenced on the Illumina HiSeq 3/4000 platform with read lengths of 150 bp. The raw data are available from the Sequence Read Archive (SRA) at NCBI (Accession ID: PRJNA715613).

Transcriptomic analysis was performed according to previous procedures^6^. For mapping of haddock RNA sequences, we chose to use Atlantic de novo cod trinity transcriptome^71^ because the official gene-model versions did not contain full length of all sequences. Of the 39426 annotated cod reference sequences, there were 34400 reference sequences with at least 10 mapped haddock reads in one sample. The RNA sequencing data were mapped to the reference sequences of the cod genes^72^ using the Bowtie aligner (RRID:SCR_005476)^73^. The mapping efficiency averaged 39%, giving an overall average of 14 million mapped paired end reads (150 bp) for each sample. Genes with less than 10 reads in both control and treatment were excluded from further analysis. Mapped reads were normalized to the total number of mapped sequences. Annotation of genes, extraction of number of mapped gene reads (Samstools idxstat), statistics (NOIseqBIO) and Kyoto Encyclopedia of Genes and Genomes (KEGG) pathway analysis was performed as described previously^6,65^. Qiagen’s Ingenuity Pathway Analysis (IPA version 01–16) was used in addition to KEGG pathway analysis. The log2(fold change) (Log2FC) and p values were extracted from the original NOISeqBIO output. Log2FC values between −1 and 1 was set to 1, as non-significant values and those below 10 reads were originally set to 0.5 and 0.75. Swiss prot IDs were combined with the Log2FC and p-value data and uploaded to IPA. This resulted in 31376 of 39426 genes (80%) mapping to the IPA database.

Figures were made using relative regulation of KEGG pathways, i.e number of genes in pathway divided by number of annotated genes in pathway. Heat maps were generated from Log2(Fold change) data in R (<u>R Core Team, 2013</u>) and Venn diagrams were created using the web-tool, Venn (http://bioinformatics.psb.ugent.be/webtools/Venn). IPA-related figures were extracted directly from the IPA app. To extract most enriched pathways (Canonical pathways and Diseases and Biofunctions) (Fig. 2) an activation z-score threshold of 3 was applied (p-value <0.05). While the p value reflects the likelihood of a pathway being significantly enriched following treatment, the z-score describes the likelihood of a directional association in our dataset. Positive and negative z-scores implies an increase or decrease in activity/activation of the enriched pathway, respectively. z-score > 2 is regarded as significant.

## Notes

### Competing Interest Statement

The authors have declared no competing interest.

## References

1 Incardona, J. P. & Scholz, N. L. The influence of heart developmental anatomy on cardiotoxicity-based adverse outcome pathways in fish. Aquat. toxicol. 177, 515–525, (2016).

2 Sørhus, E. et al. Crude oil exposures reveal roles for intracellular calcium cycling in haddock craniofacial and cardiac development. Sci. Rep. 6, 31058, (2016).

3 Perrichon, P., Donald, C. E., Sørhus, E., Harboe, T. & Meier, S. Differential developmental toxicity of crude oil in early life stages of Atlantic halibut (Hippoglossus hippoglossus). Sci. Total Environ. 770, 145349, (2021).

4 Perrichon, P. et al.. Combined effects of elevated temperature and Deepwater Horizon oil exposure on the cardiac performance of larval mahi-mahi, Coryphaena hippurus. Plos One 13, (2018).

5 Sørhus, E. et al.. Untangling mechanisms of crude oil toxicity: Linking gene expression, morphology and PAHs at two developmental stages in a cold-water fish. Sci. Total Environ. 757, 143896, (2021).

6 Sørhus, E. et al.. Novel adverse outcome pathways revealed by chemical genetics in a developing marine fish. Elife 6, (2017).

7 Lie, K. K. et al. Offshore Crude Oil Disrupts Retinoid Signaling and Eye Development in Larval Atlantic Haddock. Front. Mar. Sci. 6, (2019).

8 Cresci, A. et al.. Effects of exposure to low concentrations of oil on expression of cytochrome P4501a and routine swimming speed of Atlantic haddock (Melanogrammus aeglefinus) larvae in situ. Environ. Sci. Technol., (2020).

9 Heintz, R. A., Short, J. W. & Rice, S. D. Sensitivity of fish embryos to weathered crude oil: Part II. Increased mortality of pink salmon (Oncorhynchus gorbuscha) embryos incubating downstream from weathered Exxon Valdez crude oil. Environ. Toxicol. Chem. 18, 494–503, (1999).

10 Incardona, J. P. et al.. Very low embryonic crude oil exposures cause lasting cardiac defects in salmon and herring. Sci. Rep. 5, 13499, (2015).

11 Gardner, L. D. et al.. Cardiac remodeling in response to embryonic crude oil exposure involves unconventional NKX family members and innate immunity genes. J. Exp. Biol. 222, (2019).

12 Brette, F. et al.. Crude oil impairs cardiac excitation-contraction coupling in fish. Science 343, 772–776, (2014).

13 Brette, F. et al.. A Novel Cardiotoxic Mechanism for a Pervasive Global Pollutant. Sci. Rep. 7, 41476, (2017).

14 Ainerua, M. O. et al.. Understanding the cardiac toxicity of the anthropogenic pollutant phenanthrene on the freshwater indicator species, the brown trout (Salmo trutta): From whole heart to cardiomyocytes. Chemosphere 239, 124608, (2020).

15 Kompella, S. N., Brette, F., Hancox, J. C. & Shiels, H. A. Phenanthrene impacts zebrafish cardiomyocyte excitability by inhibiting I-Kr and shortening action potential duration. J. Gen. Physiol. 153, (2021).

16 Zhang, Y. Y., Huang, L. X., Zuo, Z. H., Chen, Y. X. & Wang, C. G. Phenanthrene exposure causes cardiac arrhythmia in embryonic zebrafish via perturbing calcium handling. Aquat. toxicol. 142, 26–32, (2013).

17 Newton, A. C., Bootman, M. D. & Scott, J. D. Second Messengers. Cold Spring Harb. Perspect. Biol. 8, (2016).

18 Marris, C. R. et al.. Polyaromatic hydrocarbons in pollution: a heart-breaking matter. J. Physiol. 598, 227–247, (2019).

19 Shwartz, Y., Farkas, Z., Stern, T., Aszodi, A. & Zelzer, E. Muscle contraction controls skeletal morphogenesis through regulation of chondrocyte convergent extension. Dev. Biol. 370, 154–163, (2012).

20 Drugbank.com. Nicardipine, 2020).

21 Koide, Y., Kimura, S., Tada, R., Kugai, N. & Yamashita, K. Activation of Glycogenolysis by the Reduction in the Extracellular Calcium-Concentration in Verapamil-Perfused Rat-Liver. Biochem. Pharmacol. 32, 517–522, (1983).

22 Mittnacht, A. J. C., London, M. J., Puskas, J. D. & Kaplan, J. A. in Kaplan’s Essentials of Cardiac Anesthesia (Second Edition) (ed Joel A. Kaplan) 322–351 (Elsevier, 2018).

23 Magnuson, J. T., Khursigara, A. J., Allmon, E. B., Esbaugh, A. J. & Roberts, A. P. Effects of Deepwater Horizon crude oil on ocular development in two estuarine fish species, red drum (Sciaenops ocellatus) and sheepshead minnow (Cyprinodon variegatus). Ecotoxicol. Environ. Saf. 166, 186–191, (2018).

24 Cvekl, A. & Wang, W.-L. Retinoic acid signaling in mammalian eye development. Exp. Eye Res. 89, 280–291, (2009).

25 Williams, A. L., Eason, J., Chawla, B. & Bohnsack, B. L. Cyp1b1 Regulates Ocular Fissure Closure Through a Retinoic Acid-Independent Pathway. Invest. Ophth. Vis. Sci. 58, (2017).

26 Berntssen, M. H. G., ørnsrud, R., Hamre, K. & Lie, K. K. Polyaromatic hydrocarbons in aquafeeds, source, effects and potential implications for vitamin status of farmed fish species: a review. Aquacult. Nutr. 21, 257–273, (2015).

27 Berntssen, M. H. G. et al. Dietary vitamin A supplementation ameliorates the effects of poly-aromatic hydrocarbons in Atlantic salmon (Salmo salar). Aquat. toxicol. 175, 171–183, (2016).

28 Incardona, J. P., Collier, T. K. & Scholz, N. L. Defects in cardiac function precede morphological abnormalities in fish embryos exposed to polycyclic aromatic hydrocarbons. Toxicol. Appl. Pharmacol. 196, 191–205, (2004).

29 Bell, M. V., Mcevoy, L. A. & Navarro, J. C. Deficit of didocosahexaenoyl phospholipid in the eyes of larval sea bass fed an essential fatty acid deficient diet. J. Fish Biol. 49, 941–952, (1996).

30 Koven, W. et al.. The effect of dietary DHA and taurine on rotifer capture success, growth, survival and vision in the larvae of Atlantic bluefin tuna (Thunnus thynnus). Aquaculture 482, 137–145, (2018).

31 Stoknes, I. S., økland, H. M., Falch, E. & Synnes, M. Fatty acid and lipid class composition in eyes and brain from teleosts and elasmobranchs. Comp. Biochem. Physiol. B Biochem. Mol. Biol. 138, 183–191, (2004).

32 Hoehne-Reitan, K. &Kjørsvik, E. Functional development of the liver and exocrine pancreas. Am. Fish. Soc. Symp. 40, 9–36, (2004).

33 Hall, T. E., Smith, P. & Johnston, I. A. Stages of embryonic development in the Atlantic cod Gadus morhua. J. Morphol. 259, 255–270, (2004).

34 Sørensen, L. et al.. Oil droplet fouling and differential toxicokinetics of polycyclic aromatic hydrocarbons in embryos of Atlantic haddock and cod. Plos One 12, e0180048, (2017).

35 Sørhus, E. et al.. Unexpected interaction with dispersed crude oil droplets drives severe toxicity in atlantic haddock embryos. Plos One 10, e0124376, (2015).

36 Xiong, K. M., Peterson, R. E. & Heideman, W. Aryl Hydrocarbon Receptor-Mediated Down-Regulation of Sox9b Causes Jaw Malformation in Zebrafish Embryos. Mol. Pharmacol. 74, 1544–1553, (2008).

37 Planchart, A. & Mattingly, C. J. 2,3,7,8-Tetrachlorodibenzo-p-dioxin upregulates FoxQ1b in zebrafish jaw primordium. Chem. Res. Toxicol. 23, 480–487, (2010).

38 Goodman, C. A., Hornberger, T. A. & Robling, A. G. Bone and skeletal muscle: Key players in mechanotransduction and potential overlapping mechanisms. Bone 80, 24–36, (2015).

39 Sato, Y. & Fujino, M. Inhibition of Excitation-Contraction Coupling by a Ca Channel Blocker Nicardipine at Low-Temperature in Frog Twitch Fibers. Jpn. J. Physiol. 37, 93–108, (1987).

40 Knight, R. D. & Schilling, T. F. Cranial neural crest and development of the head skeleton. Adv. Exp. Med. Biol. 589, 120–133, (2006).

41 Stifani, N. Motor neurons and the generation of spinal motor neuron diversity. Cell Neurosci. 8, 1–22, (2014).

42 Saccà, S. C., Cutolo, C. A., Ferrari, D., Corazza, P. & Traverso, C. E. The Eye, Oxidative Damage and Polyunsaturated Fatty Acids. Nutrients 10, 668, (2018).

43 Gorusupudi, A., Liu, A., Hageman, G. S. & Bernstein, P. S. Associations of human retinal very long-chain polyunsaturated fatty acids with dietary lipid biomarkers. J. Lipid Res. 57, 499–508, (2016).

44 Morais, S. et al.. Molecular and Functional Characterization of Elovl4 Genes in Sparus aurata and Solea senegalensis Pointing to a Critical Role in Very Long-Chain (>C(24)) Fatty Acid Synthesis during Early Neural Development of Fish. Int. J. Mol. Sci. 21, (2020).

45 Serrano, R. et al.. Identification of new, very long-chain polyunsaturated fatty acids in fish by gas chromatography coupled to quadrupole/time-of-flight mass spectrometry with atmospheric pressure chemical ionization. Analytical and Bioanalytical Chemistry 413, 1039–1046, (2021).

46 Agbaga, M. P., Mandal, M. N. & Anderson, R. E. Retinal very long-chain PUFAs: new insights from studies on ELOVL4 protein. J. Lipid Res. 51, 1624–1642, (2010).

47 Zampelas, A. & Magriplis, E. New Insights into Cholesterol Functions: A Friend or an Enemy? Nutrients 11, (2019).

48 Wallaert, C. & Babin, P. J. Thermal adaptation affects the fatty acid composition of plasma phospholipids in trout. Lipids 29, 373–376, (1994).

49 Broniatowski, M., Binczycka, M., Wojcik, A., Flasinski, M. & Wydro, P. Polycyclic aromatic hydrocarbons in model bacterial membranes - Langmuir monolayer studies. Bba-Biomembranes 1859, 2402–2412, (2017).

50 McGruer, V. et al.. Effects of Phenanthrene Exposure on Cholesterol Homeostasis and Cardiotoxicity in Zebrafish Embryos. Environ. Toxicol. Chem., (2021).

51 Roderick, H. L., Berridge, M. J. & Bootman, M. D. Calcium-induced calcium release. Curr. Biol. 13, R425, (2003).

52 Tanito, M. et al.. High levels of retinal membrane docosahexaenoic acid increase susceptibility to stress-induced degeneration. J. Lipid Res. 50, 807–819, (2009).

53 Matsuzaka, T. & Shimano, H. Elovl6: a new player in fatty acid metabolism and insulin sensitivity. J. Mol. Med. 87, 379–384, (2009).

54 Ralston, J. C., Matravadia, S., Gaudio, N., Holloway, G. P. & Mutch, D. M. Polyunsaturated Fatty Acid Regulation of Adipocyte FADS1 and FADS2 Expression and Function. Obesity 23, 725–728, (2015).

55 Kamler, E. Resource allocation in yolk-feeding fish. Rev. Fish Biol. Fisher. 18, 143–200, (2008).

56 Ross, S. A., McCaffery, P. J., Drager, U. C. & De Luca, L. M. Retinoids in embryonal development. Physiol. Rev. 80, 1021–1054, (2000).

57 Alsop, D. H., Brown, S. B. & van der Kraak, G. J. Dietary retinoic acid induces hindlimb and eye deformities in Xenopus laevis. Environ. Sci. Technol. 38, 6290–6299, (2004).

58 Le, H. G., Dowling, J. E. & Cameron, D. J. Early retinoic acid deprivation in developing zebrafish results in microphthalmia. Vis. Neurosci. 29, 219–228, (2012).

59 Yamamoto, Y., Zolfaghari, R. & Ross, A. C. Regulation of CYP26 (cytochrome P450RAI) mRNA expression and retinoic acid metabolism by retinoids and dietary vitamin A in liver of mice and rats. Faseb J. 14, 2119–2127, (2000).

60 Murphy, K. A., Quadro, L. & White, L. A. The intersection between the aryl hydrocarbon receptor (AhR)-and retinoic acid-signaling pathways. Vitam. Horm. 75, 33–67, (2007).

61 Chambers, D., Wilson, L., Maden, M. & Lumsden, A. RALDH-independent generation of retinoic acid during vertebrate embryogenesis by CYP1B1. Development 134, 1369–1383, (2007).

62 Hawkins, S. A., Billiard, S. M., Tabash, S. P., Brown, R. S. & Hodson, P. V. Altering cytochrome P4501A activity affects polycyclic aromatic hydrocarbon metabolism and toxicity in rainbow trout (Oncorhynchus mykiss). Environ. Toxicol. Chem. 21, 1845–1853, (2002).

63 Terakita, A. The opsins. Genome Biol. 6, 213, (2005).

64 Burns, M. E. & Pugh, E. N., Jr. RGS9 concentration matters in rod phototransduction. Biophys. J. 97, 1538–1547, (2009).

65 Sørhus, E. et al.. Developmental transcriptomics in Atlantic haddock: Illuminating pattern formation and organogenesis in non-model vertebrates. Dev. Biol. 411, 301–313, (2016).

66 Vergauwen, L. et al.. A high throughput passive dosing format for the Fish Embryo Acute Toxicity test. Chemosphere 139, 9–17, (2015).

67 Sørensen, L. et al.. Establishing a link between composition and toxicity of offshore produced waters using comprehensive analysis techniques - A way forward for discharge monitoring? Sci. Total Environ. 694, 133682, (2019).

68 Sørensen, L., Meier, S. &Mjøs, S. A. Application of gas chromatography/tandem mass spectrometry to determine a wide range of petrogenic alkylated polycyclic aromatic hydrocarbons in biotic samples. Rapid Commun Mass Sp 30, 2052–2058, (2016).

69 Meier, S., Mjos, S. A., Joensen, H. & Grahl-Nielsen, O. Validation of a one-step extraction/methylation method for determination of fatty acids and cholesterol in marine tissues. J. Chromatogr. A 1104, 291–298, (2006).

70 Wasta, Z. &Mjøs, S. A. A database of chromatographic properties and mass spectra of fatty acid methyl esters from omega-3 products. J. Chromatogr. A 1299, 94–102, (2013).

71 Hanna, E. M. et al.. ReCodLiver0.9: Overcoming Challenges in Genome-Scale Metabolic Reconstruction of a Non-model Species. Front. Mol. Biosci. 7, 591406, (2020).

72 Star, B. et al.. The genome sequence of Atlantic cod reveals a unique immune system. Nature 477, 207–210, (2011).

73 Langmead, B., Trapnell, C., Pop, M. & Salzberg, S. L. Ultrafast and memory-efficient alignment of short DNA sequences to the human genome. Genome Biol. 10, (2009).

