## Supplementary Information for "Cardiac dysfunction affects eye development and vision by reducing supply of lipids in fish"

#### Supplementary figures:

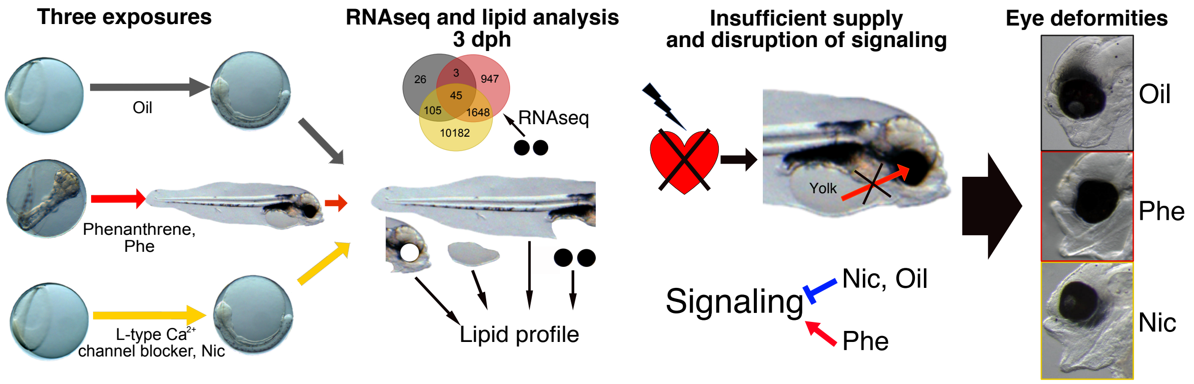


**Figure S1: Graphical abstract.** Graphical representation of the findings in the study

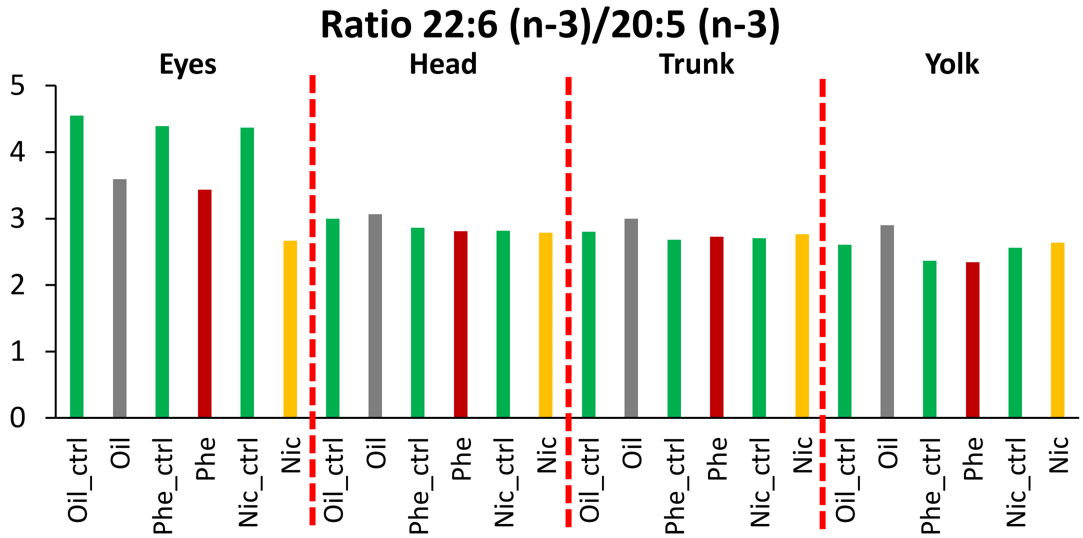


**Figure S2: Ratios between 22:6 (n-3)/20:5 (n-3) in eyes, head, trunk and yolk**. Oil exposure; Oil, Oil experiment control; Oil_ctr, Phenanthrene exposure; Phe, Phenanthrene experiment control; Phe_ctr, Nicardipine hydrochloride exposure; Nic; Nicardipine experiment control; Nic_ctr.

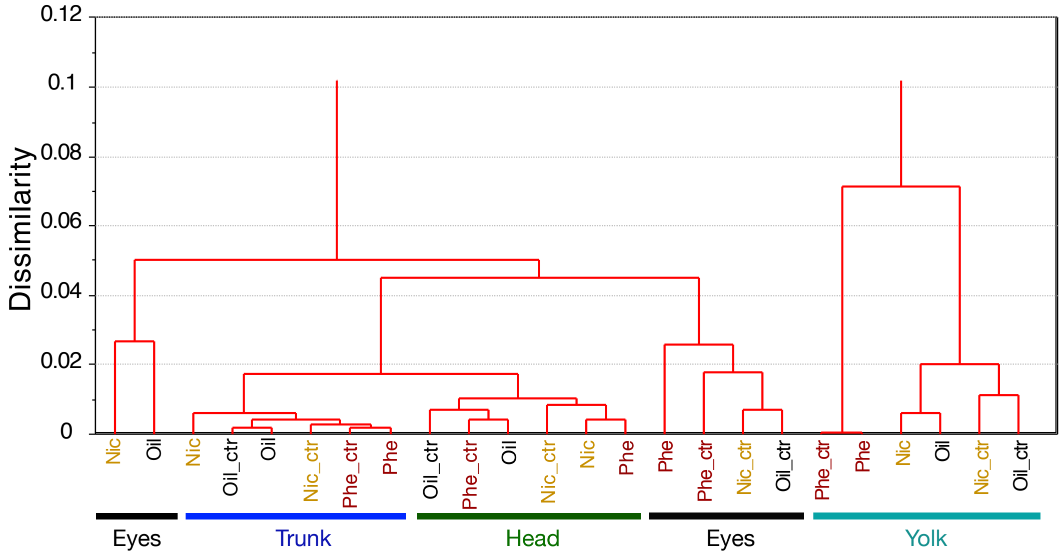


**Figure S3. Classification of control and treatment samples from eyes, head, trunk and yolk**. Oil exposure; Oil, Oil experiment control; Oil_ctr, Phenanthrene exposure; Phe, Phenanthrene experiment control; Phe_ctr, Nicardipine hydrochloride exposure; Nic; Nicardipine experiment control; Nic_ctr.

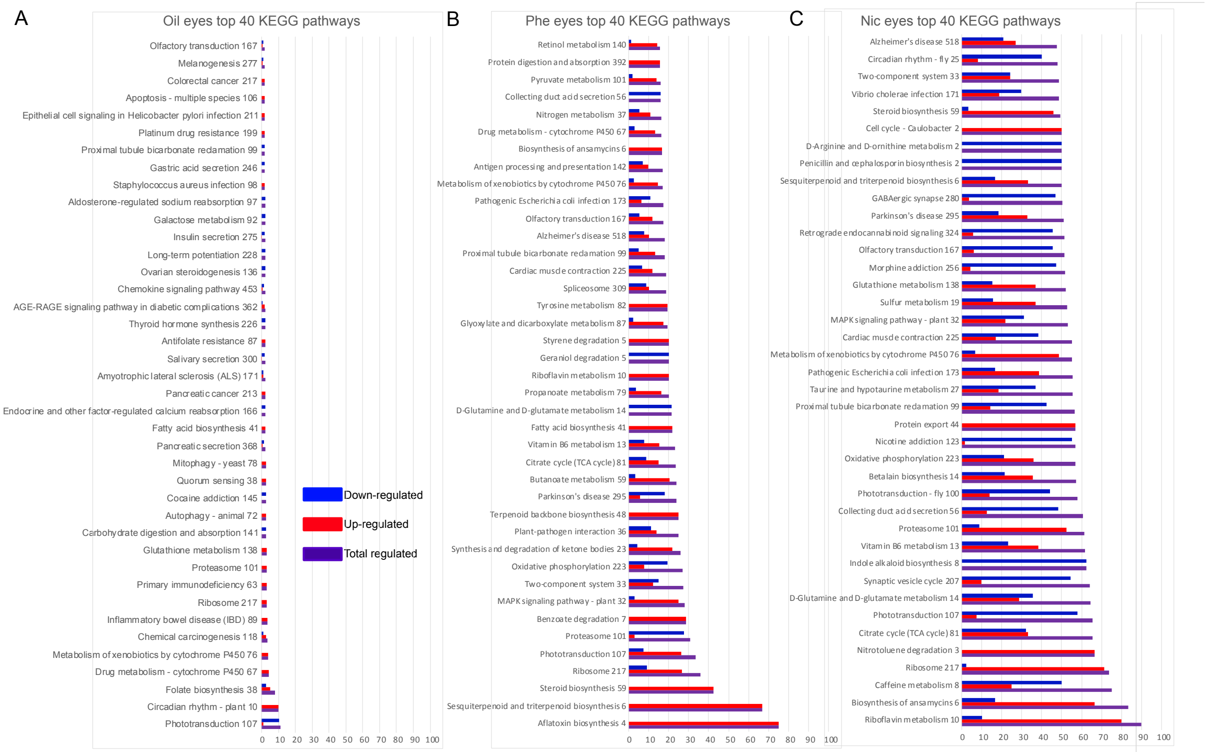


**Supplementary Figure S4: Top 40 relative KEGG pathways.** Threshold for fold change and p-value was set to 1.5 and <0.05, respectively. Number of genes regulated in pathways was divided by number of total annotated genes pathway and are presented as percentage on x-axis. Number behind the Pathway name represent number of annotated genes in respective pathway.

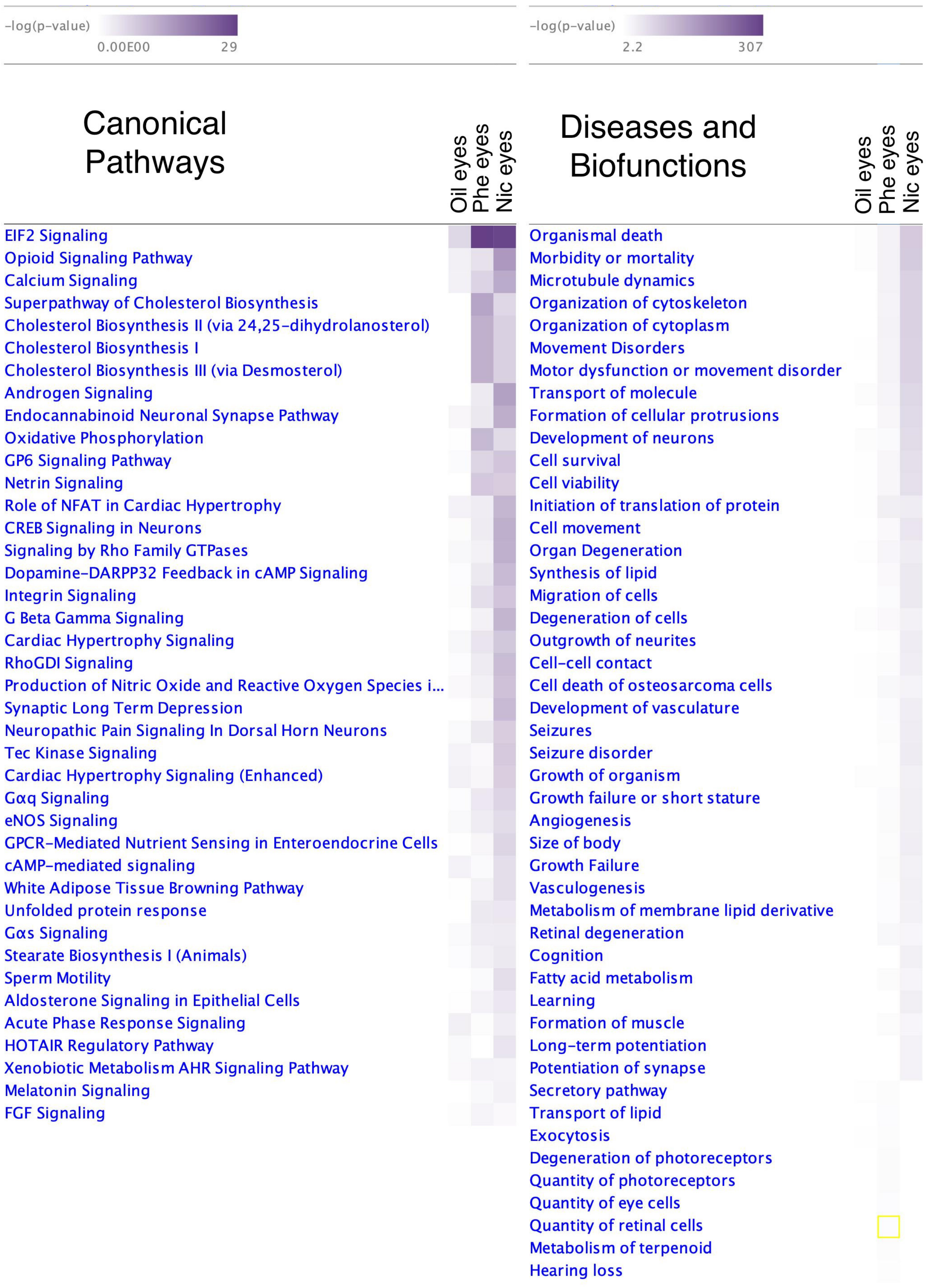


**Supplementary figure S5: IPA Pathways.** Canonical pathways (A) and Disease and Biofunctions (B) displayed with p-value. Threshold for z-score and p-value was set to 3 and <0.05 respectively.

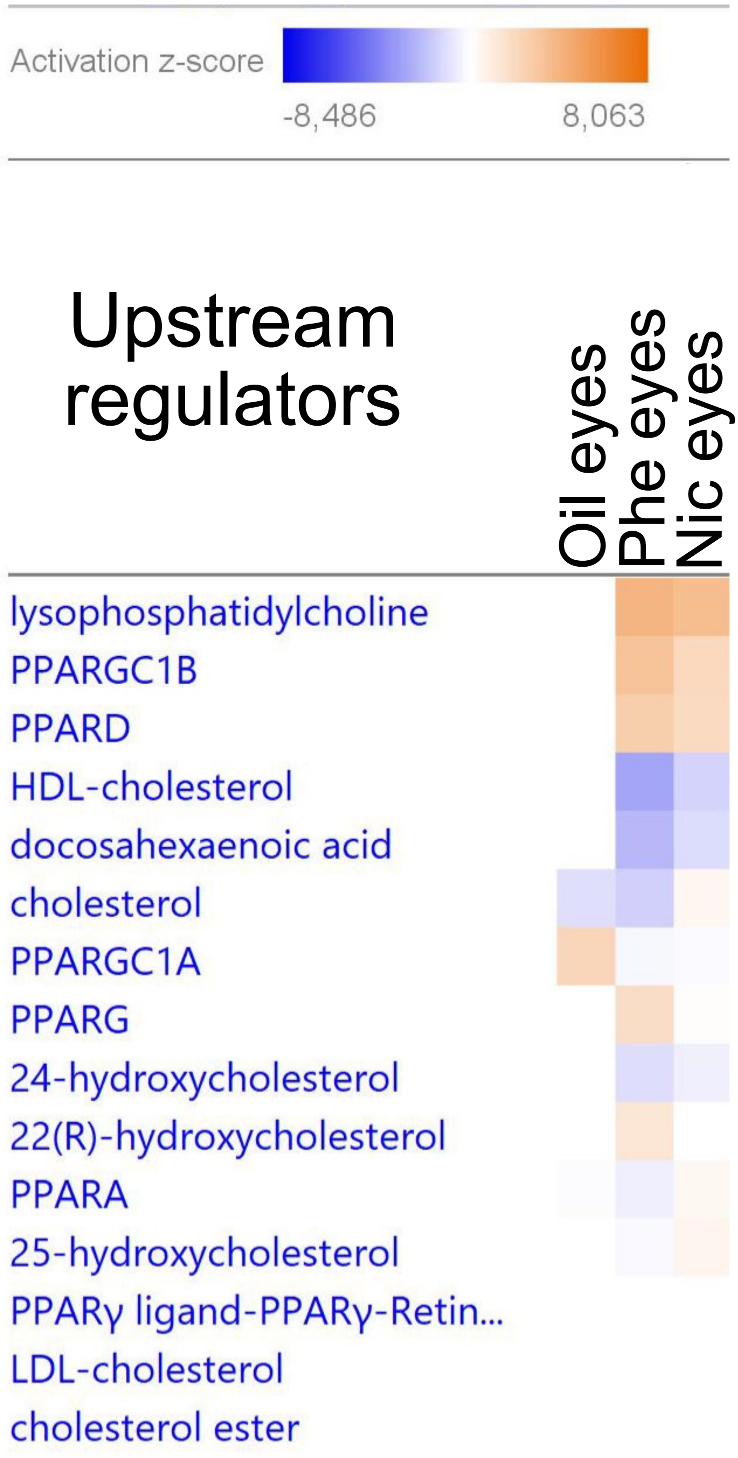


**Supplementary Figure S6. Up-stream regulators identified by IPA analysis.**

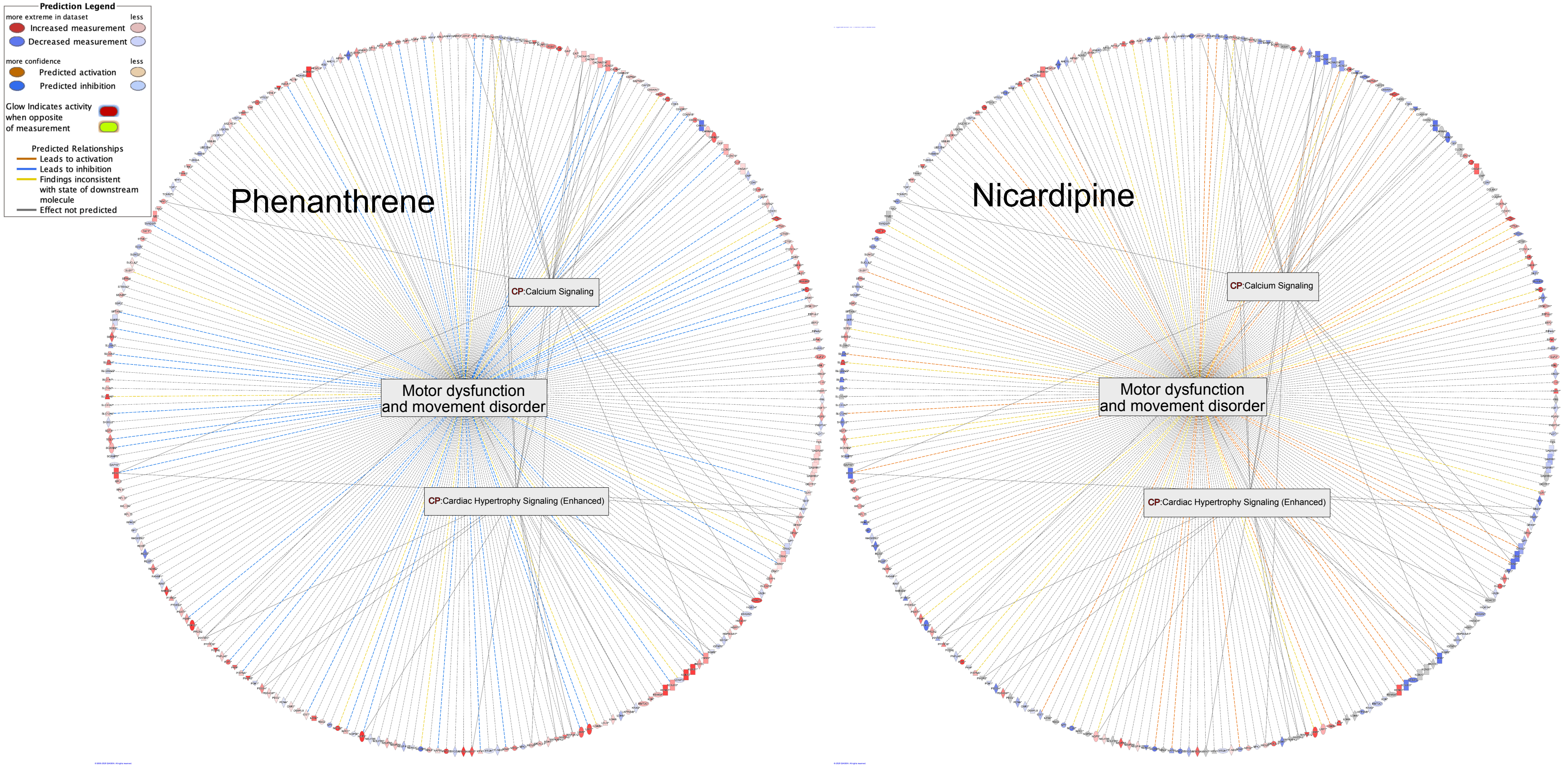


**Supplementary Figure S7: Motor function and movement disorder.** IPA Motor dysfunction and movement disorder gene wheel. Visual web of genes in Motor dysfunction and movement disorder overlapping with the Canonical pathways (CP), Calcium signaling (21 molecules) and Cardiac hypertrophy signaling (Enhanced) (19 molecules).

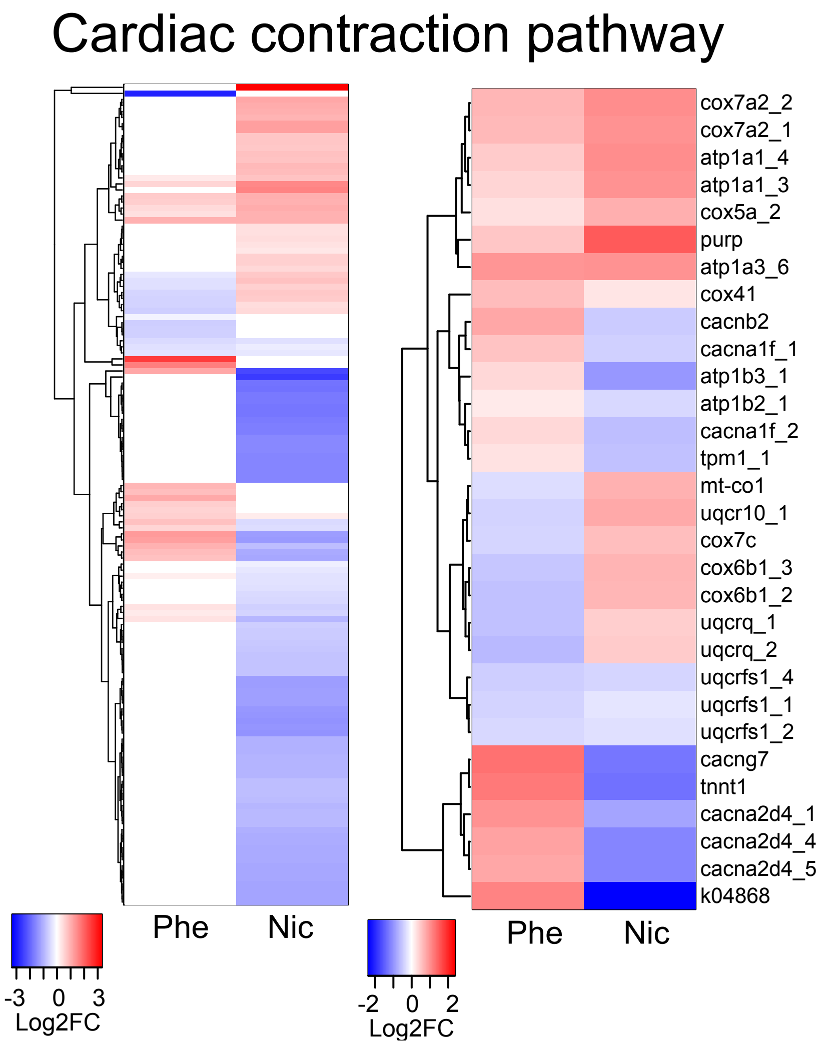


**Supplementary Figure S8**: Cardiac contraction pathway as an indicator for muscular function. KEGG Cardiac contraction pathway. Left column: All regulated in genes in Phe and/or Nic eyes. Right column: Regulated genes in both Phe and Nic eyes.

### Supplementary tables:

**Supplementary Table 1**. Phenanthrene water concentration (μg Phe/L). After 1 day of exposure the water concentration decreased and was steady on the level until exposure stop.

| PAH | Average, day 0 | Averages, day 1-3 | Percent decrease after day 0 | Average water conc (μg/L) in period +/-SD |
|---|---|---|---|---|
| PHE | 351 | 149 | 57% | 196 +/-14.2 |

**Supplementary Table 2.** Fatty acids and cholesterol amount and fatty acids profiles from eyes, head, trunk and yolk from control and Oil treatment. The analysis was performed on pooled samples from 10 larvae. FA; fatty acids, Chol; cholesterol, SFA; saturated fatty acids, MUFA; monounsaturated fatty acids, PUFA; polyunsaturated fatty acids.

| | Control Eyes | Oil Eyes | Control Head | Oil Head | Control Trunk | Oil Trunk | Control Yolk | Oil Yolk |
|---|---|---|---|---|---|---|---|---|
| FA (µg/larvae) | 0.79 | 0.28 | 2.93 | 1.37 | 2.47 | 2.24 | 1.66 | 1.82 |
| Chol (µg/larvae) | 0.12 | 0.04 | 0.51 | 0.23 | 0.35 | 0.31 | 0.07 | 0.08 |
| FA profile (% of total Fas) | | | | | | | | |
| 14:0 | 0.62 | 1.12 | 0.69 | 0.76 | 0.83 | 0.85 | 0.82 | 0.73 |
| Iso 15:0 | 0.02 | 0.01 | 0.03 | 0.04 | 0.05 | 0.04 | 0.09 | 0.08 |
| 15:0 | 0.20 | 0.21 | 0.20 | 0.19 | 0.21 | 0.21 | 0.24 | 0.23 |
| Iso 16:0 | 0.01 | 0.00 | 0.02 | 0.02 | 0.03 | 0.03 | 0.04 | 0.04 |
| 16:0 | 22.49 | 26.46 | 20.48 | 20.97 | 19.20 | 19.47 | 15.71 | 15.77 |
| Iso 17:0 | 0.17 | 0.12 | 0.21 | 0.21 | 0.27 | 0.20 | 0.18 | 0.18 |
| Antiso 17:0 | 0.06 | 0.07 | 0.05 | 0.06 | 0.06 | 0.05 | 0.06 | 0.07 |
| 17:0 | 0.27 | 0.32 | 0.26 | 0.26 | 0.26 | 0.27 | 0.26 | 0.26 |
| 18:0 | 7.22 | 9.59 | 6.11 | 6.30 | 5.44 | 5.39 | 6.37 | 4.91 |
| 20:0 | 0.09 | 0.14 | 0.09 | 0.10 | 0.08 | 0.09 | 0.06 | 0.04 |
| **∑SFA** | **31.15** | **38.04** | **28.13** | **28.90** | **26.43** | **26.60** | **23.83** | **22.32** |
| 16:1 (n-11) | 0.08 | 0.09 | 0.14 | 0.14 | 0.16 | 0.14 | 0.06 | 0.07 |
| 16:1 (n-9) | 1.28 | 1.82 | 1.83 | 2.06 | 1.94 | 1.96 | 0.61 | 0.80 |
| 16:1 (n-7) | 0.99 | 1.58 | 1.29 | 1.44 | 1.56 | 1.59 | 1.28 | 1.17 |
| 16:1 (n-5) | 0.06 | 0.07 | 0.08 | 0.07 | 0.08 | 0.09 | 0.11 | 0.09 |
| 17:1 (n-8) | 0.13 | 0.18 | 0.20 | 0.15 | 0.21 | 0.22 | 0.18 | 0.19 |
| 16:1 n-10, 7Me | 0.30 | 0.41 | 0.35 | 0.35 | 0.40 | 0.34 | 0.29 | 0.28 |
| 18:1 (n-11) | 0.14 | 0.09 | 0.10 | 0.25 | 0.20 | 0.15 | 0.18 | 0.29 |
| 18:1 (n-9) | 6.88 | 8.61 | 10.42 | 10.67 | 11.25 | 11.63 | 14.00 | 14.06 |
| 18:1 (n-7) | 2.59 | 2.83 | 3.38 | 3.59 | 3.51 | 3.72 | 4.18 | 4.63 |
| 18:1 (n-5) | 0.10 | 0.10 | 0.15 | 0.15 | 0.17 | 0.17 | 0.11 | 0.13 |
| 20:1 (n-11) | 0.01 | 0.03 | 0.04 | 0.02 | 0.05 | 0.06 | 0.14 | 0.11 |
| 20:1 (n-9) | 0.72 | 0.90 | 1.27 | 1.46 | 1.39 | 1.51 | 1.40 | 1.21 |
| 20:1 (n-7) | 0.05 | 0.04 | 0.06 | 0.07 | 0.06 | 0.07 | 0.07 | 0.06 |
| 22:1 (n-11) | 0.09 | 0.09 | 0.12 | 0.11 | 0.09 | 0.06 | 0.19 | 0.15 |
| 22:1 (n-9) | 0.05 | 0.13 | 0.05 | 0.06 | 0.08 | 0.08 | 0.11 | 0.13 |
| 22:1 (n-7) | 0.03 | 0.04 | 0.06 | 0.04 | 0.03 | 0.03 | 0.02 | 0.02 |
| 24:1 (n-9) | 0.41 | 0.57 | 0.60 | 0.52 | 0.63 | 0.61 | 2.15 | 1.82 |
| **∑MUFA** | **13.96** | **17.65** | **20.21** | **21.19** | **21.88** | **22.51** | **25.14** | **25.27** |
| 16:4 (n-1) | 0.00 | 0.01 | 0.04 | 0.02 | 0.04 | 0.04 | 0.02 | 0.02 |
| 18:4 (n-1) | 0.00 | 0.00 | 0.00 | 0.04 | 0.00 | 0.05 | 0.00 | 0.00 |
| 16:2 (n-4) | 0.01 | 0.01 | 0.04 | 0.04 | 0.06 | 0.06 | 0.06 | 0.05 |
| 16:3 (n-4) | 0.00 | 0.01 | 0.01 | 0.01 | 0.02 | 0.02 | 0.02 | 0.02 |
| 18:2 (n-4) | 0.10 | 0.09 | 0.15 | 0.14 | 0.18 | 0.18 | 0.15 | 0.15 |
| 16:2 (n-6) | 0.08 | 0.09 | 0.12 | 0.12 | 0.15 | 0.14 | 0.05 | 0.07 |
| 18:2 (n-6) | 1.67 | 1.93 | 2.55 | 2.61 | 3.19 | 3.16 | 3.52 | 3.51 |
| 18:3 (n-6) | 0.01 | 0.04 | 0.03 | 0.04 | 0.04 | 0.04 | 0.03 | 0.02 |
| 20:2 (n-6) | 0.34 | 0.39 | 0.43 | 0.47 | 0.43 | 0.44 | 0.57 | 0.47 |
| 20:3 (n-6) | 0.07 | 0.06 | 0.11 | 0.09 | 0.12 | 0.10 | 0.13 | 0.11 |
| 20:4 (n-6) | 1.32 | 1.27 | 1.57 | 1.47 | 1.51 | 1.39 | 4.01 | 3.63 |
| 22:4 (n-6) | 0.09 | 0.04 | 0.19 | 0.11 | 0.15 | 0.14 | 0.14 | 0.14 |
| 22:5 (n-6) | 0.22 | 0.19 | 0.27 | 0.27 | 0.29 | 0.26 | 0.52 | 0.45 |
| 18:3 (n-3) | 0.18 | 0.19 | 0.29 | 0.28 | 0.38 | 0.35 | 0.26 | 0.25 |
| 18:4 (n-3) | 0.21 | 0.24 | 0.70 | 0.67 | 1.03 | 0.91 | 0.62 | 0.71 |
| 20:3 (n-3) | 0.05 | 0.05 | 0.07 | 0.06 | 0.09 | 0.07 | 0.10 | 0.09 |
| 20:4 (n-3) | 0.22 | 0.17 | 0.09 | 0.24 | 0.09 | 0.28 | 0.03 | 0.05 |
| 20:5 (n-3) | 8.71 | 8.26 | 10.76 | 10.17 | 11.06 | 10.35 | 10.61 | 10.28 |
| 21:5 (n-3) | 0.07 | 0.06 | 0.11 | 0.10 | 0.12 | 0.11 | 0.11 | 0.10 |
| 22:5 (n-3) | 1.52 | 1.37 | 1.57 | 1.50 | 1.49 | 1.48 | 2.04 | 2.12 |
| 22:6 (n-3) | 39.65 | 29.67 | 32.20 | 31.13 | 30.94 | 30.98 | 27.63 | 29.77 |
| 24:5 (n-3) | 0.33 | 0.13 | 0.28 | 0.26 | 0.24 | 0.25 | 0.22 | 0.23 |
| 24:6 (n-3) | 0.06 | 0.06 | 0.07 | 0.08 | 0.08 | 0.08 | 0.17 | 0.18 |
| **∑PUFA** | **54.89** | **44.31** | **51.66** | **49.91** | **51.69** | **50.89** | **51.03** | **52.42** |
| ∑PUFA (n-6) | 3.79 | 4.01 | 5.28 | 5.18 | 5.88 | 5.67 | 8.98 | 8.40 |
| ∑PUFA (n-3) | 51.00 | 40.18 | 46.15 | 44.48 | 45.52 | 44.87 | 41.80 | 43.78 |
| ∑Chol | 14.62 | 15.05 | 17.51 | 16.97 | 14.02 | 13.76 | 4.28 | 4.32 |

**Supplementary Table 3.** Fatty acids and cholesterol amount and fatty acids profiles from eyes, head, trunk and yolk from control and phenanthrene treatment. The analysis was performed on pooled samples from 10 larvae. FA; fatty acids, Chol; cholesterol, SFA; saturated fatty acids, MUFA; monounsaturated fatty acids, PUFA; polyunsaturated fatty acids.

| | Control | Phe | Control | Phe | Control | Phe | Control | Phe |
|---|---|---|---|---|---|---|---|---|
| | Eyes | Eyes | Head | Head | Trunk | Trunk | Yolk | Yolk |
| FA (µg/larvae) | 0.84 | 0.60 | 2.32 | 2.32 | 2.78 | 2.70 | 0.68 | 0.60 |
| Chol (µg/larvae) | 0.12 | 0.09 | 0.42 | 0.40 | 0.43 | 0.39 | 0.04 | 0.03 |
| FA profile (% of total Fas) | | | | | | | | |
| 14:0 | 0.68 | 0.85 | 0.73 | 0.79 | 0.81 | 0.85 | 0.88 | 0.84 |
| Iso 15:0 | 0.03 | 0.03 | 0.04 | 0.04 | 0.04 | 0.05 | 0.09 | 0.10 |
| 15:0 | 0.18 | 0.20 | 0.22 | 0.19 | 0.21 | 0.21 | 0.26 | 0.25 |
| Iso 16:0 | 0.02 | 0.02 | 0.03 | 0.03 | 0.03 | 0.03 | 0.05 | 0.05 |
| 16:0 | 20.53 | 22.91 | 20.59 | 20.78 | 19.16 | 18.88 | 13.37 | 13.27 |
| Iso 17:0 | 0.16 | 0.14 | 0.24 | 0.24 | 0.21 | 0.22 | 0.15 | 0.11 |
| Antiso 17:0 | 0.07 | 0.08 | 0.07 | 0.06 | 0.07 | 0.06 | 0.09 | 0.08 |
| 17:0 | 0.28 | 0.27 | 0.27 | 0.26 | 0.27 | 0.26 | 0.35 | 0.34 |
| 18:0 | 6.96 | 7.24 | 6.38 | 6.37 | 5.57 | 5.44 | 9.90 | 9.93 |
| 20:0 | 0.08 | 0.10 | 0.09 | 0.09 | 0.08 | 0.08 | 0.08 | 0.09 |
| **∑SFA** | **28.99** | **31.85** | **28.66** | **28.85** | **26.46** | **26.09** | **25.23** | **25.06** |
| 16:1 (n-11) | 0.10 | 0.13 | 0.14 | 0.18 | 0.15 | 0.16 | 0.03 | 0.05 |
| 16:1 (n-9) | 1.01 | 1.48 | 1.71 | 1.82 | 1.80 | 1.75 | 0.52 | 0.51 |
| 16:1 (n-7) | 0.86 | 1.36 | 1.34 | 1.56 | 1.61 | 1.71 | 0.96 | 0.94 |
| 16:1 (n-5) | 0.06 | 0.08 | 0.08 | 0.09 | 0.10 | 0.12 | 0.06 | 0.08 |
| 17:1 (n-8) | 0.16 | 0.20 | 0.23 | 0.23 | 0.24 | 0.23 | 0.16 | 0.17 |
| 16:1 n-10, 7Me | 0.26 | 0.34 | 0.35 | 0.34 | 0.37 | 0.34 | 0.28 | 0.25 |
| 18:1 (n-11) | 0.17 | 0.16 | 0.14 | 0.25 | 0.12 | 0.17 | 0.12 | 0.14 |
| 18:1 (n-9) | 8.05 | 8.79 | 11.26 | 11.15 | 11.87 | 12.00 | 12.19 | 12.14 |
| 18:1 (n-7) | 2.96 | 2.95 | 3.58 | 3.53 | 3.79 | 3.76 | 4.41 | 4.41 |
| 18:1 (n-5) | 0.09 | 0.12 | 0.15 | 0.16 | 0.17 | 0.18 | 0.08 | 0.09 |
| 20:1 (n-11) | 0.07 | 0.01 | 0.06 | 0.05 | 0.06 | 0.06 | 0.09 | 0.15 |
| 20:1 (n-9) | 0.87 | 0.91 | 1.29 | 1.28 | 1.39 | 1.41 | 1.80 | 1.83 |
| 20:1 (n-7) | 0.05 | 0.04 | 0.06 | 0.06 | 0.05 | 0.08 | 0.08 | 0.07 |
| 22:1 (n-11) | 0.15 | 0.12 | 0.14 | 0.12 | 0.12 | 0.12 | 0.29 | 0.33 |
| 22:1 (n-9) | 0.13 | 0.09 | 0.09 | 0.06 | 0.13 | 0.11 | 0.23 | 0.24 |
| 22:1 (n-7) | 0.14 | 0.01 | 0.05 | 0.04 | 0.03 | 0.04 | 0.08 | 0.10 |
| 24:1 (n-9) | 0.83 | 0.47 | 0.55 | 0.62 | 0.63 | 0.65 | 3.68 | 3.73 |
| **∑MUFA** | **15.98** | **17.26** | **21.26** | **21.61** | **22.65** | **22.95** | **25.10** | **25.26** |
| 16:4 (n-1) | 0.07 | 0.04 | 0.04 | 0.04 | 0.06 | 0.06 | 0.02 | 0.03 |
| 18:4 (n-1) | 0.00 | 0.03 | 0.04 | 0.00 | 0.06 | 0.06 | 0.01 | 0.02 |
| 16:2 (n-4) | 0.01 | 0.02 | 0.04 | 0.05 | 0.06 | 0.06 | 0.03 | 0.03 |
| 16:3 (n-4) | 0.00 | 0.01 | 0.03 | 0.02 | 0.03 | 0.03 | 0.00 | 0.01 |
| 18:2 (n-4) | 0.10 | 0.12 | 0.15 | 0.15 | 0.20 | 0.19 | 0.08 | 0.12 |
| 16:2 (n-6) | 0.07 | 0.09 | 0.12 | 0.15 | 0.16 | 0.14 | 0.04 | 0.03 |
| 18:2 (n-6) | 1.85 | 2.15 | 2.64 | 2.66 | 3.23 | 3.25 | 2.75 | 2.78 |
| 18:3 (n-6) | 0.02 | 0.04 | 0.04 | 0.04 | 0.05 | 0.04 | 0.02 | 0.02 |
| 20:2 (n-6) | 0.35 | 0.42 | 0.43 | 0.46 | 0.42 | 0.49 | 0.64 | 0.66 |
| 20:3 (n-6) | 0.10 | 0.08 | 0.10 | 0.12 | 0.12 | 0.14 | 0.16 | 0.19 |
| 20:4 (n-6) | 1.96 | 1.37 | 1.56 | 1.50 | 1.52 | 1.46 | 7.19 | 7.14 |
| 22:4 (n-6) | 0.09 | 0.09 | 0.19 | 0.26 | 0.19 | 0.20 | 0.15 | 0.14 |
| 22:5 (n-6) | 0.28 | 0.23 | 0.26 | 0.27 | 0.27 | 0.29 | 0.44 | 0.48 |
| 18:3 (n-3) | 0.22 | 0.27 | 0.28 | 0.31 | 0.35 | 0.39 | 0.14 | 0.15 |
| 18:4 (n-3) | 0.18 | 0.17 | 0.52 | 0.84 | 0.83 | 0.81 | 0.49 | 0.49 |
| 20:3 (n-3) | 0.06 | 0.07 | 0.07 | 0.07 | 0.07 | 0.08 | 0.09 | 0.11 |
| 20:4 (n-3) | 0.18 | 0.23 | 0.24 | 0.27 | 0.29 | 0.32 | 0.13 | 0.14 |
| 20:5 (n-3) | 8.79 | 9.77 | 10.69 | 10.58 | 11.16 | 11.02 | 10.33 | 10.37 |
| 21:5 (n-3) | 0.08 | 0.09 | 0.11 | 0.11 | 0.12 | 0.13 | 0.08 | 0.08 |
| 22:5 (n-3) | 1.61 | 1.65 | 1.53 | 1.56 | 1.54 | 1.53 | 2.10 | 2.12 |
| 22:6 (n-3) | 38.57 | 33.55 | 30.61 | 29.73 | 29.84 | 29.98 | 24.40 | 24.30 |
| 24:5 (n-3) | 0.31 | 0.28 | 0.29 | 0.26 | 0.24 | 0.25 | 0.32 | 0.21 |
| 24:6 (n-3) | 0.11 | 0.12 | 0.09 | 0.08 | 0.08 | 0.07 | 0.07 | 0.05 |
| **∑PUFA** | **55.03** | **50.89** | **50.08** | **49.54** | **50.89** | **50.96** | **49.68** | **49.68** |
| ∑PUFA (n-6) | 4.72 | 4.47 | 5.34 | 5.47 | 5.96 | 6.01 | 11.38 | 11.46 |
| ∑PUFA (n-3) | 50.12 | 46.21 | 44.44 | 43.80 | 44.52 | 44.56 | 38.15 | 38.01 |
| ∑Chol | 14.32 | 15.03 | 18.20 | 17.09 | 15.30 | 14.41 | 5.20 | 5.17 |

**Supplementary Table 4.** Fatty acids and cholesterol amount and fatty acids profiles from eyes, head, trunk and yolk from control and nicardipine hydrochloride treatment. The analysis was performed on pooled samples from 10 larvae. FA; fatty acids, Chol; cholesterol, SFA; saturated fatty acids, MUFA; monounsaturated fatty acids, PUFA; polyunsaturated fatty acids.

| | Control Eyes | Nic Eyes | Control Head | Nic Head | Control Trunk | Nic Trunk | Control Yolk | Nic Yolk |
|---|---|---|---|---|---|---|---|---|
| FA (µg/larvae) | 0.66 | 0.45 | 3.05 | 1.95 | 2.93 | 3.15 | 1.63 | 2.01 |
| Chol (µg/larvae) | 0.10 | 0.07 | 0.54 | 0.34 | 0.48 | 0.49 | 0.09 | 0.11 |
| FA profile (% of total Fas) | | | | | | | | |
| 14:0 | 0.75 | 1.24 | 0.99 | 0.87 | 0.86 | 0.92 | 0.70 | 0.71 |
| Iso 15:0 | 0.02 | 0.04 | 0.04 | 0.04 | 0.04 | 0.05 | 0.07 | 0.07 |
| 15:0 | 0.22 | 0.25 | 0.35 | 0.19 | 0.22 | 0.22 | 0.21 | 0.22 |
| Iso 16:0 | 0.02 | 0.03 | 0.04 | 0.03 | 0.03 | 0.04 | 0.04 | 0.04 |
| 16:0 | 22.82 | 25.12 | 21.24 | 19.73 | 19.17 | 19.58 | 14.09 | 15.68 |
| Iso 17:0 | 0.15 | 0.13 | 0.25 | 0.20 | 0.21 | 0.23 | 0.18 | 0.19 |
| Antiso 17:0 | 0.07 | 0.10 | 0.08 | 0.07 | 0.07 | 0.07 | 0.06 | 0.06 |
| 17:0 | 0.30 | 0.24 | 0.31 | 0.22 | 0.27 | 0.27 | 0.28 | 0.28 |
| 18:0 | 7.24 | 9.53 | 6.39 | 5.95 | 5.11 | 4.96 | 5.64 | 4.45 |
| 20:0 | 0.09 | 0.14 | 0.11 | 0.10 | 0.08 | 0.09 | 0.05 | 0.03 |
| **∑SFA** | **31.68** | **36.83** | **29.80** | **27.39** | **26.07** | **26.41** | **21.32** | **21.73** |
| 16:1 (n-11) | 0.09 | 0.12 | 0.15 | 0.16 | 0.16 | 0.14 | 0.06 | 0.05 |
| 16:1 (n-9) | 1.21 | 1.92 | 2.28 | 2.05 | 1.75 | 1.81 | 0.60 | 0.67 |
| 16:1 (n-7) | 0.96 | 2.08 | 1.39 | 1.83 | 1.70 | 1.87 | 1.34 | 1.39 |
| 16:1 (n-5) | 0.07 | 0.07 | 0.09 | 0.10 | 0.10 | 0.09 | 0.08 | 0.10 |
| 17:1 (n-8) | 0.18 | 0.24 | 0.22 | 0.23 | 0.23 | 0.25 | 0.19 | 0.21 |
| 16:1 n-10, 7Me | 0.29 | 0.39 | 0.35 | 0.31 | 0.37 | 0.31 | 0.27 | 0.25 |
| 18:1 (n-11) | 0.11 | 0.11 | 0.20 | 0.13 | 0.12 | 0.12 | 0.16 | 0.28 |
| 18:1 (n-9) | 7.55 | 11.08 | 10.87 | 11.24 | 11.83 | 12.51 | 13.81 | 14.06 |
| 18:1 (n-7) | 2.74 | 3.13 | 3.43 | 3.48 | 3.65 | 3.72 | 4.50 | 4.35 |
| 18:1 (n-5) | 0.11 | 0.13 | 0.19 | 0.15 | 0.17 | 0.18 | 0.13 | 0.12 |
| 20:1 (n-11) | 0.06 | 0.03 | 0.06 | 0.06 | 0.07 | 0.06 | 0.14 | 0.11 |
| 20:1 (n-9) | 0.75 | 1.02 | 1.17 | 1.49 | 1.34 | 1.48 | 1.53 | 1.19 |
| 20:1 (n-7) | 0.04 | 0.06 | 0.06 | 0.06 | 0.07 | 0.08 | 0.08 | 0.06 |
| 22:1 (n-11) | 0.10 | 0.15 | 0.13 | 0.14 | 0.08 | 0.13 | 0.24 | 0.22 |
| 22:1 (n-9) | 0.24 | 0.13 | 0.41 | 0.09 | 0.11 | 0.11 | 0.14 | 0.10 |
| 22:1 (n-7) | 0.09 | 0.06 | 0.07 | 0.05 | 0.03 | 0.04 | 0.07 | 0.05 |
| 24:1 (n-9) | 0.45 | 0.61 | 0.52 | 0.61 | 0.65 | 0.58 | 2.20 | 1.54 |
| **∑MUFA** | **15.06** | **21.36** | **21.63** | **22.25** | **22.45** | **23.53** | **25.60** | **24.79** |
| 16:4 (n-1) | 0.03 | 0.03 | 0.05 | 0.06 | 0.07 | 0.08 | 0.03 | 0.03 |
| 18:4 (n-1) | 0.02 | 0.03 | 0.04 | 0.05 | 0.06 | 0.06 | 0.03 | 0.04 |
| 16:2 (n-4) | 0.01 | 0.03 | 0.04 | 0.05 | 0.07 | 0.07 | 0.05 | 0.05 |
| 16:3 (n-4) | 0.01 | 0.02 | 0.03 | 0.02 | 0.03 | 0.05 | 0.02 | 0.03 |
| 18:2 (n-4) | 0.10 | 0.11 | 0.16 | 0.16 | 0.18 | 0.20 | 0.14 | 0.17 |
| 16:2 (n-6) | 0.09 | 0.10 | 0.14 | 0.15 | 0.14 | 0.13 | 0.05 | 0.05 |
| 18:2 (n-6) | 1.84 | 2.45 | 2.60 | 2.78 | 3.28 | 3.29 | 3.43 | 3.48 |
| 18:3 (n-6) | 0.01 | 0.04 | 0.05 | 0.05 | 0.05 | 0.05 | 0.02 | 0.04 |
| 20:2 (n-6) | 0.30 | 0.31 | 0.41 | 0.44 | 0.41 | 0.43 | 0.59 | 0.48 |
| 20:3 (n-6) | 0.07 | 0.08 | 0.09 | 0.12 | 0.12 | 0.12 | 0.16 | 0.13 |
| 20:4 (n-6) | 1.34 | 1.46 | 1.42 | 1.43 | 1.46 | 1.28 | 4.55 | 3.51 |
| 22:4 (n-6) | 0.07 | 0.08 | 0.18 | 0.28 | 0.22 | 0.19 | 0.19 | 0.14 |
| 22:5 (n-6) | 0.26 | 0.20 | 0.25 | 0.26 | 0.30 | 0.25 | 0.54 | 0.43 |
| 18:3 (n-3) | 0.26 | 0.30 | 0.32 | 0.33 | 0.40 | 0.44 | 0.24 | 0.29 |
| 18:4 (n-3) | 0.14 | 0.27 | 0.55 | 0.62 | 0.83 | 0.69 | 0.47 | 0.49 |
| 20:3 (n-3) | 0.06 | 0.05 | 0.06 | 0.07 | 0.08 | 0.08 | 0.11 | 0.10 |
| 20:4 (n-3) | 0.21 | 0.19 | 0.25 | 0.27 | 0.31 | 0.33 | 0.18 | 0.23 |
| 20:5 (n-3) | 8.67 | 9.36 | 10.48 | 10.88 | 11.20 | 10.73 | 11.18 | 11.35 |
| 21:5 (n-3) | 0.08 | 0.09 | 0.10 | 0.11 | 0.13 | 0.13 | 0.10 | 0.14 |
| 22:5 (n-3) | 1.43 | 1.39 | 1.48 | 1.58 | 1.51 | 1.48 | 2.13 | 2.08 |
| 22:6 (n-3) | 37.85 | 24.92 | 29.49 | 30.30 | 30.28 | 29.66 | 28.57 | 29.93 |
| 24:5 (n-3) | 0.27 | 0.18 | 0.27 | 0.26 | 0.26 | 0.25 | 0.24 | 0.25 |
| 24:6 (n-3) | 0.11 | 0.08 | 0.11 | 0.10 | 0.08 | 0.06 | 0.06 | 0.04 |
| **∑PUFA** | **53.25** | **41.81** | **48.57** | **50.36** | **51.48** | **50.06** | **53.08** | **53.47** |
| ∑PUFA (n-6) | 3.98 | 4.74 | 5.13 | 5.51 | 5.98 | 5.74 | 9.54 | 8.25 |
| ∑PUFA (n-3) | 49.10 | 36.85 | 43.12 | 44.52 | 45.08 | 43.85 | 43.27 | 44.90 |
| ∑Chol | 14.51 | 14.69 | 17.55 | 17.27 | 16.52 | 15.61 | 5.28 | 5.56 |

**Supplementary Table 5:** Normalized total fatty acids and cholesterol in the eyes. TFA/diam,Total fatty acids (µg/larva)/diameter of the eye; Chol/diam, Cholesterol (µg/larva)/diameter of the eye. C, control; Phe, phenanthrene; Nic, Nicardipine

| | TFA/diam | Chol/diam |
|---|---|---|
| | **Eyes** | **Eyes** |
| **Control** | 0.00269 | 0.00041 |
| **Oil** | 0.00106 | 0.00016 |
| **Oil/C (%)** | 39% | 39% |
| **Control** | 0.00312 | 0.00045 |
| **Phe** | 0.00240 | 0.00036 |
| **Phe/C (%)** | 77% | 81% |
| **Control** | 0.00203 | 0.00031 |
| **Nic** | 0.00174 | 0.00025 |
| **Nic/C (%)** | 86% | 83% |

**Supplementary Data:**

**Supplementary Data 1**: Concentration of Polycyclic aromatic hydrocarbons (PAHs) in water in the Oil Exposure. Sum of total PAHs and individual PAHs and sum of groups of individual PAHs. C0; non alkylated, C1-C4; alkylated homologues BT: Benzothiophene, NAP: Naphtalene, FLU: Fluorene, PHE: Phenanthrene, DBT: Dibenzothiophene, PYR: Pyrene, CHR: Chrysene

**Supplementary Data 2:** Concentration of Polycyclic aromatic hydrocarbons (PAHs) in tissue in the Oil Exposure. Sum of total PAHs and individual PAHs and sum of groups of individual PAHs. C0; non alkylated, C1-C4; alkylated homologues BT: Benzothiophene, NAP: Naphtalene, FLU: Fluorene, PHE: Phenanthrene, DBT: Dibenzothiophene, PYR: Pyrene, CHR: Chrysene.

**Supplementary Data 3**: Differentially expressed genes in KEGG fat metabolism and PPAR signaling. Differentially expressed genes (p<0.05) in either Phe-eyes or Nic-eyes in fat metabolism and PPAR signaling. KEGG; Kyoto Encyclopedia of Genes and Genomes, IMR_gene_ID; The exclusive gene identification after mapping at Institute of Marine Research, Phe; Phenanthrene eyes, Nic; Nicardipine eyes, C; Control, FC; Fold change, Oil_eye; Oil treatment experiment, Phe_eye; Phe treatment experiment, Nic_eye; Nic treatment experiment, mean; mean transcripts, swissprot; Swiss-Prot annotation, prob; probability (p-value = 1-prob).

**Supplementary Data 4:** Genes in KEGG Calcium signaling pathway. Differentially expressed genes (p<0.05) in either Phe-eyes or Nic-eyes in Calcium signaling pathway. Phe; Phenanthrene eyes, Nic; Nicardipine eyes, FC; Fold change.

**Supplementary Data 5**: Genes in KEGG Cardiac contraction pathway. Differentially expressed genes ($p<0.05$) in either Phe-eyes or Nic-eyes in Cardiac contraction pathway. Phe; Phenanthrene eyes, Nic; Nicardipine eyes, FC; Fold change.
